# Multi-sample Full-length Transcriptome Analysis of 22 Breast Cancer Clinical Specimens with Long-Read Sequencing

**DOI:** 10.1101/2020.07.15.199851

**Authors:** Shinichi Namba, Toshihide Ueno, Shinya Kojima, Yosuke Tanaka, Satoshi Inoue, Fumishi Kishigami, Noriko Maeda, Tomoko Ogawa, Shoichi Hazama, Yuichi Shiraishi, Hiroyuki Mano, Masahito Kawazu

## Abstract

Although transcriptome alteration is considered as one of the essential drivers of carcinogenesis, conventional short-read RNAseq technology has limited researchers from directly exploring full-length transcripts, only focusing on individual splice sites. We developed a pipeline for Multi-Sample long-read Transcriptome Assembly, MuSTA, and showed through simulations that it enables construction of transcriptome from the transcripts expressed in target samples and more accurate evaluation of transcript usage. We applied it to 22 breast cancer clinical specimens to successfully acquire cohort-wide full-length transcriptome from long-read RNAseq data. By comparing isoform existence and expression between estrogen receptor positive and triple-negative subtypes, we obtained a comprehensive set of subtype-specific isoforms and differentially used isoforms which consisted of both known and unannotated isoforms. We have also found that exon-intron structure of fusion transcripts tends to depend on their genomic regions, and have found three-piece fusion transcripts that were transcribed from complex structural rearrangements. For example, a three-piece fusion transcript resulted in aberrant expression of an endogenous retroviral gene, *ERVFRD-1*, which is normally expressed exclusively in placenta and supposed to protect fetus from maternal rejection, and expression of which were increased in several TCGA samples with *ERVFRD-1* fusions. Our analyses of real clinical specimens and simulated data provide direct evidence that full-length transcript sequencing in multiple samples can add to our understanding of cancer biology and genomics in general.

## Introduction

Transcriptome is an important determinant of cellular phenotype^1^, and its alteration is a major driver of oncogenesis as well as DNA alteration^2^. In some cases, aberrant splicing regulation are observed recurrently^3^ and considered as independent drivers from somatic mutations^4^. It has been recognized that some genes have cancer-specific splicing isoforms which are the driving forces of cancer proliferation, such as *PKM2* in Warburg effect^5^, long non-coding RNA *PNUTS* in epithelial-mesenchymal transition^6^, and *BRAF* exon3-9 in chemo-resistance^7^. Therefore, exploring splicing patterns in cancer is indispensable for further understanding tumorigenesis as well as other hallmarks of cancer.

Some groups have recently conducted comprehensive researches of cancer-specific alternative splicing using the data from the Cancer Genome Atlas^8–10^ and International Cancer Genome Consortium^2^, and found that RNA alteration affects cancer genes complementary to DNA alteration^2^. However, all these studies depend on RNAseq technology, with the drawback in that RNAseq produces relatively short read fragments and needs imputation with de Bruijn graph to achieve full-length transcripts. This is why previous analyses were limited to individual splice site abnormalities and could neither directly nor efficiently target consequent transcripts. It is especially difficult to quantify the gene expression in transcript level, and annotation list with incomplete set of isoforms results in insufficient estimation accuracy^11^. Transcript expression shows cell-type specific pattern^12^, and there are far more isoforms than those transcripts registered on reference annotation^13^. Therefore, unless we use complete reference transcripts of target cells, it is difficult to correctly evaluate the transcripts usage.

Single-molecule real-time (SMRT) sequencing technology^14^ can sequence far longer reads of 10 kbp and over, and its recent advent enabled us to read the full-length of transcripts without fragmentation. Although raw reads have relatively high error rate (~ 10%) of base calling, it can be compensated for by clustering into highly accurate consensus sequences (IsoSeq protocol)^15^ and hybrid error correction with RNAseq data^16,17^. Several groups have used this technology to capture high resolution transcriptomes of eukaryotes^18–20^ including human^13^, most of which have successfully discovered unannotated or mis-annotated transcripts. These revealed their transcriptome diversity and previously undescribed transcript regulations, such as exon inclusion/skipping coupling at distal cassette exons^21,22^ and splicing coupling with transcription initiation^23^.

However, this sequencing has been utilized for obtaining cancer transcriptome in only a few studies^24,25^, and has yet to be applied to neither clinical specimens nor more than a few individual samples. The main reason is that there is no suitable software that yields transcriptome through long-read RNA sequencing from multiple samples. Even in one sample level, although long-read technologies obtain consensus sequences from multiple erroneous reads, there are many redundant consensus sequences; they tend to have shared genomic structure when mapped to the reference genome and are distinguished only by sequencing errors. Therefore, it was difficult to compare the existence or expression of transcripts between samples, preventing us from surveying cancer transcriptome at cancer-type or subtype wide level.

In this report, we first developed a pipeline for Multi-Sample long-read Transcriptome Assembly, MuSTA, and evaluated its detection ability of differential transcript usage (DTU) by using a SMRT transcriptome simulator, simlady. We then sequenced 22 breast cancer clinical specimens in total, of which fourteen were triple-negative breast cancer (TNBC) and eight were estrogen receptor or progesterone receptor positive (ER-positive) breast cancer, to acquire comprehensive full-length transcriptome of these two subtypes of breast cancer.

76.5% of the isoforms were previously unannotated ones, and subtype-specific or differently used isoforms not only contained isoforms from known subtype-specific genes, but novel transcripts as well. Furthermore, we’ve detected relationships between exon-intron structure of fusion transcripts and their genomic regions, and found three-piece fusion transcripts that transcribed from three distinct genomic regions involved in complex structural alterations.

## Results

### Overview of methods

We created a new algorithm to combine IsoSeq cluster reads from multiple samples (Fig. 1). Here, by merging IsoSeq reads which were considered to be derived from one transcript (Methods), we obtained a catalog of non-redundant transcripts according to each sample, and subsequently merged them in all samples, followed by SQANTI^26^ filtering, which removes potential artifact transcripts by a random forest algorithm, eventually creating cohort-wide transcriptome. Furthermore, this enabled us to compare the number of uniquely associated full-length non-chimeric (FLNC) reads (hereafter referred as PBcount) between samples. In addition, alignment of short-reads to the created transcriptome enables us to estimate gene expression just as conventional analyses, and to compare group wide expression such as differential gene expression (DGE) and DTU. As a pipeline with all these algorithms, we named the pipeline as “MuSTA”, Multi-Sample long-read Transcriptome Assembly.

**Figure 1.**
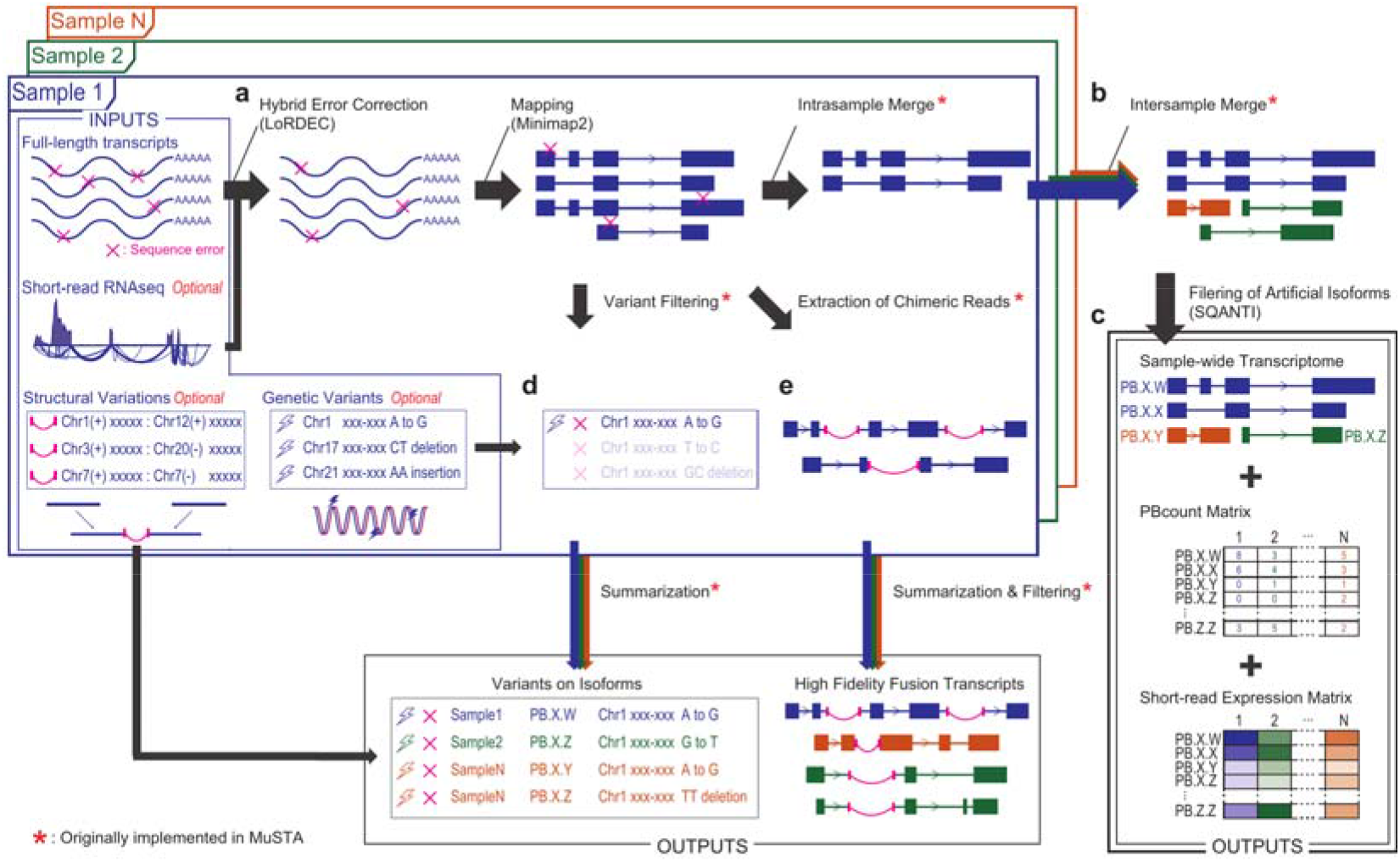
A schematic view of MuSTA workflow. **a-c**, First, IsoSeq cluster reads go through hybrid error correction if the user specifies; otherwise, this step will be skipped (**a**). Then, those reads are mapped to a reference genome with minimap2. The reads mapped to a single genomic region are merged into non-redundant isoforms in two steps, in intra-sample (**a**) and inter-sample (**b**) manners, consecutively. Short-read RNAseq data is aligned to the isoforms, and transcript per million (TPM) is calculated for the usage in SQANTI filter (**c**). SQANTI classifies the merged isoforms by comparing with a reference transcript annotation, and exclude artificial isoforms with a random forest algorithm. In each sample, and for each original cluster read, the number of full-length non-chimeric (FLNC) reads is summed up. This process was limited to when the original cluster read was linked to only one isoform. The number of FLNC reads yielded was named as “PBcount.” Then, short-read TPM is again calculated for the SQANTI-passed isoforms. **d**, The original reads contain sequence mismatches against the genome, and these mismatches are also linked to the merged isoforms. The mismatches can be filtered with user-specified genomic variants. Finally, the SQANTI-passed isoforms are reported with the information about PBcount, short-read TPM, and sequence mismatches of original reads. **e**, As for the reads which are separately mapped to multiple genomic regions, their genomic positions and mismatches against the reference genome are summarized. Of these chimeric reads, transcripts consisted of splicing junctions from the reference transcript annotation or SQANTI-passed isoforms are selected and reported. Furthermore, given the structural variation data, MuSTA categorizes transcripts according to whether there are associated structural variations. All of the procedures using short-read RNAseq data or variation data are optional, and users can execute MuSTA as long as they have IsoSeq reads.

### Cohort-wide transcriptome enables more accurate inference of transcript usage

DGE shows the variability of expression in gene level. On the other hand, variability in the proportion of isoforms in transcript level is called DTU. We evaluated the performance of MuSTA for DTU detection. Although there’re multiple long-read genomic sequence generators for simulation such as PBsim^27^, SimLoRD^28^, and FASTQsim^29^, none are designed for cDNA sequencing. Therefore, we’ve created a simulator for long-read RNA sequencing, simlady (SIMulator for Long read transcriptome Analysis with RNA DecaY model, Supplementary Fig. 1 and Methods), with the aid of the sequencing error model implemented in SimLoRD.

By using this, we’ve generated several long-read and short-read RNAseq simulation data sets where random genes were assigned to those with DTU (Methods). These data were analyzed with MuSTA, and the detection performance of DTU was evaluated under several conditions (Supplementary Figs. 2-4, Supplementary Notes).

Next, we’ve conducted a more realistic simulation based on the full-spliced match (FSM) and novel in catalog (NIC) isoforms in the breast cancer dataset described in the next section (Methods). FSM and NIC are the categories of isoforms described in ref. 26, where FSM isoforms are the isoforms of which splice junctions are completely matched to known isoforms; NIC isoforms contain at least one novel splicing junctions, however, they consist of known splicing donors and acceptors. We permutated the log-averaged expression of FSM isoforms and NIC isoforms separately, and randomly set DGE and DTU. We tested five conditions with NIC ratio against all DTU isoforms of 0, 0.25, 0.5, 0.75, and 1. Even under the value of 0 (i.e. all DTU transcripts were FSM), we observed higher precision and compatible recall for DTU inference with the MuSTA-derived annotation than GENCODE (Fig. 2). As the NIC rate increased, the MuSTA-derived annotation showed stable precision and recall, while these with GENCODE for transcript-level DTU inference decreased to 0. Although we note that transcripts can be more diverse in real situations than this simulation where the transcripts were restricted to the FSM and NIC isoforms from the breast cancer dataset, this result showed the importance of constructing transcriptome from the transcripts expressed in target samples.

**Figure 2.**
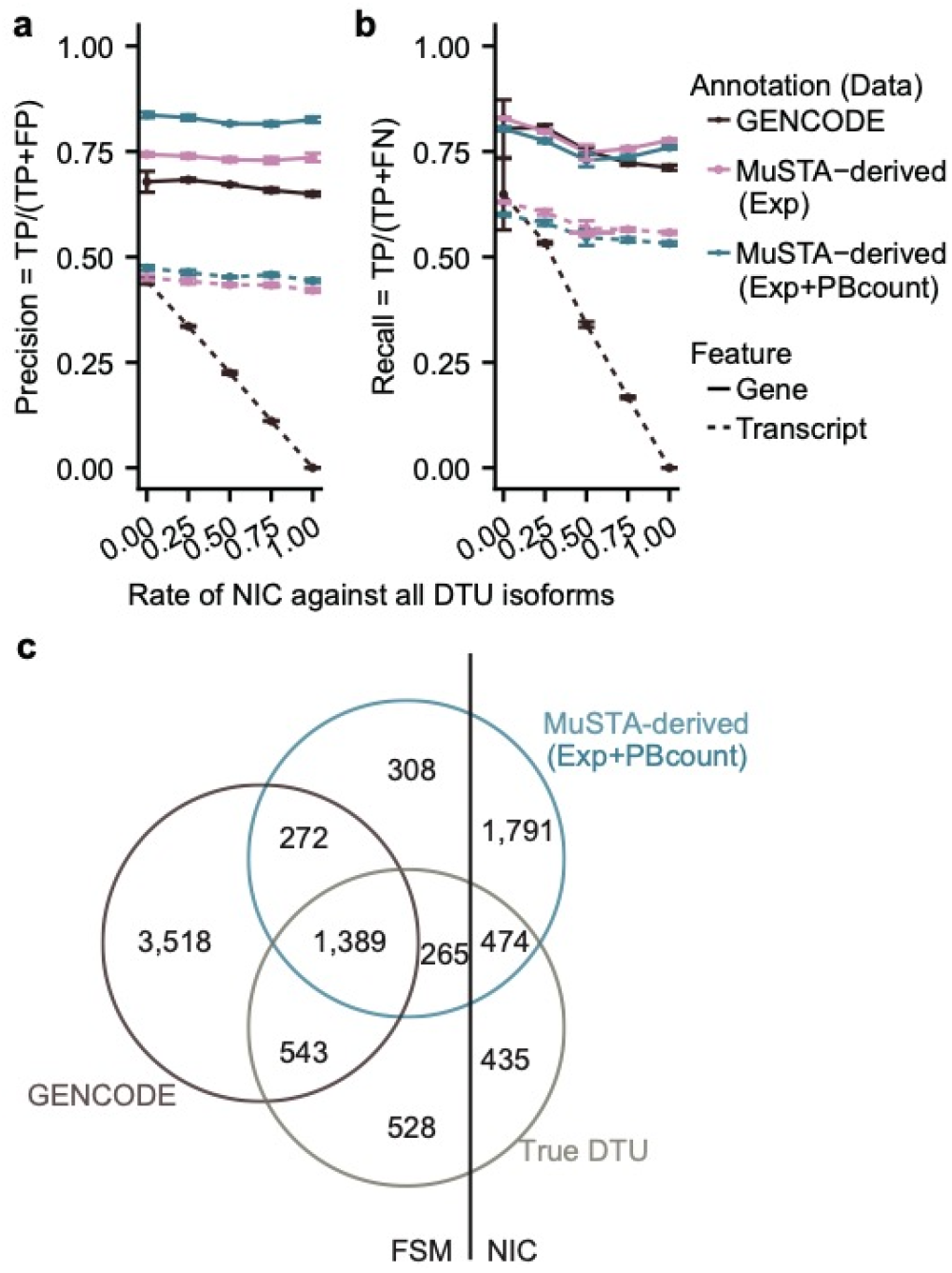
Simulations of differential transcript usage (DTU) inference with different novel in catalog (NIC) rates against all DTU isoforms. **a** and **b**, The precision (**a**) and the recall rate (**b**) of DTU inference using either GENCODE or the MuSTA-derived transcriptome. The dots represent the mean and the error bars represent the standard error of three independent simulations. **c**, A Venn diagram of DTU isoforms for a representative simulation with NIC rate of 0.25. Exp, expression; TP, true positive; FP, false positive; FN, false negative.

### Application to 22 breast cancer clinical specimens

Breast cancer is one of the most prevalent types of cancer and is typically classified into three categories, ER-positive, HER2 (erbB-2) positive, and TNBC. TNBC includes those with BRCAness, which is characterized by *BRCA1*, *BRCA2*, or *RAD51C* inactivation and consequent homologous recombination deficiency^30–32^. Breast cancer of TNBC and BRCAness have shown distinct mutational signatures compared with other types of breast cancer^31–35^. Though they have also been distinguished at the point of splice site abnormalities^10^, their detailed splicing characteristics are largely unknown. While ER-positive and HER2-positive breast cancer can be treated by hormone therapy and tyrosine-kinase inhibitor respectively, there are few specific medication strategies for TNBC, except for Poly ADP-ribose polymerase (PARP) inhibitors for BRCA-mutated cancer at clinical trials^36^. In order to simplify the comparison, we used two subtypes of ER-positive breast cancer and TNBC, and RNA samples from eight ER-positive breast cancer and fourteen TNBC clinical specimens were subjected to IsoSeq in order to obtain MuSTA transcriptome of these samples (Methods and Supplementary Fig. 5).

Number of consensus reads was between 180,179 to 523,638 reads (median 263,378); number of FLNC reads was between 159,121 to 397,067 reads (median 232,131), (which fell in the long-read depth in the simulation dataset); number of polished clusters was between 93,672 to 207,017 reads (median 124,292). By using MuSTA, 818,620 non-redundant isoforms were obtained. There were 344,504 isoforms that passed SQANTI, including 263,711 (76.5%) unannotated isoforms. Among those isoforms, 344,429 isoforms were mapped on autosomes or chromosome X, and of those, 288,674 isoforms had multiple exons. The number of SQANTI-passed isoforms detected in each sample was between 29,246 to 58,756 (median 39,313) (Supplementary Table 1, 2).

We identified 3081 unannotated multi-exonic genes. Most of them were detected only in one sample, but 41 were detected in multiple samples. Of these multi-exonic genes, ten genes were unannotated in GENCODE v28, which we used throughout this paper, but newly annotated in GENCODE v34. Eight out of the ten genes were detected in one sample. Furthermore, MuSTA-transcriptome covered 17/115 translated but unannotated open reading frames constructed in ref. 37. These results suggested that MuSTA-transcriptome successfully captured isoforms which were actually present but unannotated. In addition, because SQANTI doesn’t acknowledge genomic variants, transcripts with splice site created by mutation will be deleted as artifacts. Since the number of such isoforms were limited, as we could only find 7 of these isoforms across all the samples, we considered this problem as having a trivial or at most minor effects (Supplementary Table 3).

Relationship between short-read expression, long-read detection, and PBcount were investigated in Supplementary Notes and Supplementary Fig. 6. We also examined alternative splicing in the transcriptome and found a large number of mutually exclusive exons, and molecularly association / molecularly and mutually exclusivity between distal exons (Supplementary Fig. 7 and Supplementary Notes).

We observed strong heterogeneity of detected transcripts between samples even in the same subtypes, where more than half of the isoforms were detected only in one sample (Fig. 3a). Further, the number of detected isoforms decreased as the number of samples that generated isoforms increased up to 19 samples, indicating that the majority of isoforms were not ubiquitous. On the other hand, when the number of samples exceeded 19, the number of isoforms increased (Fig. 3a), suggesting that these isoforms are ubiquitous and essential house-keeping transcripts. Next, in order to see whether we used the sufficient number of samples, we incremented the number of analyzed samples one by one and applied MuSTA (Fig. 3b). Although the graph didn’t reach a plateau due to the aforementioned heterogeneity, we were able to yield a consistent number of isoforms that were detected in more than 80% of the samples, indicating that we were able to successfully detect most of the essential transcripts, while larger cohort is required for the thorough investigation of heterogeneity of transcriptome. These data indicated the biphasic distribution of isoforms with strong heterogeneity of minor isoforms and the ubiquitous existence of essential transcripts.

**Figure 3.**
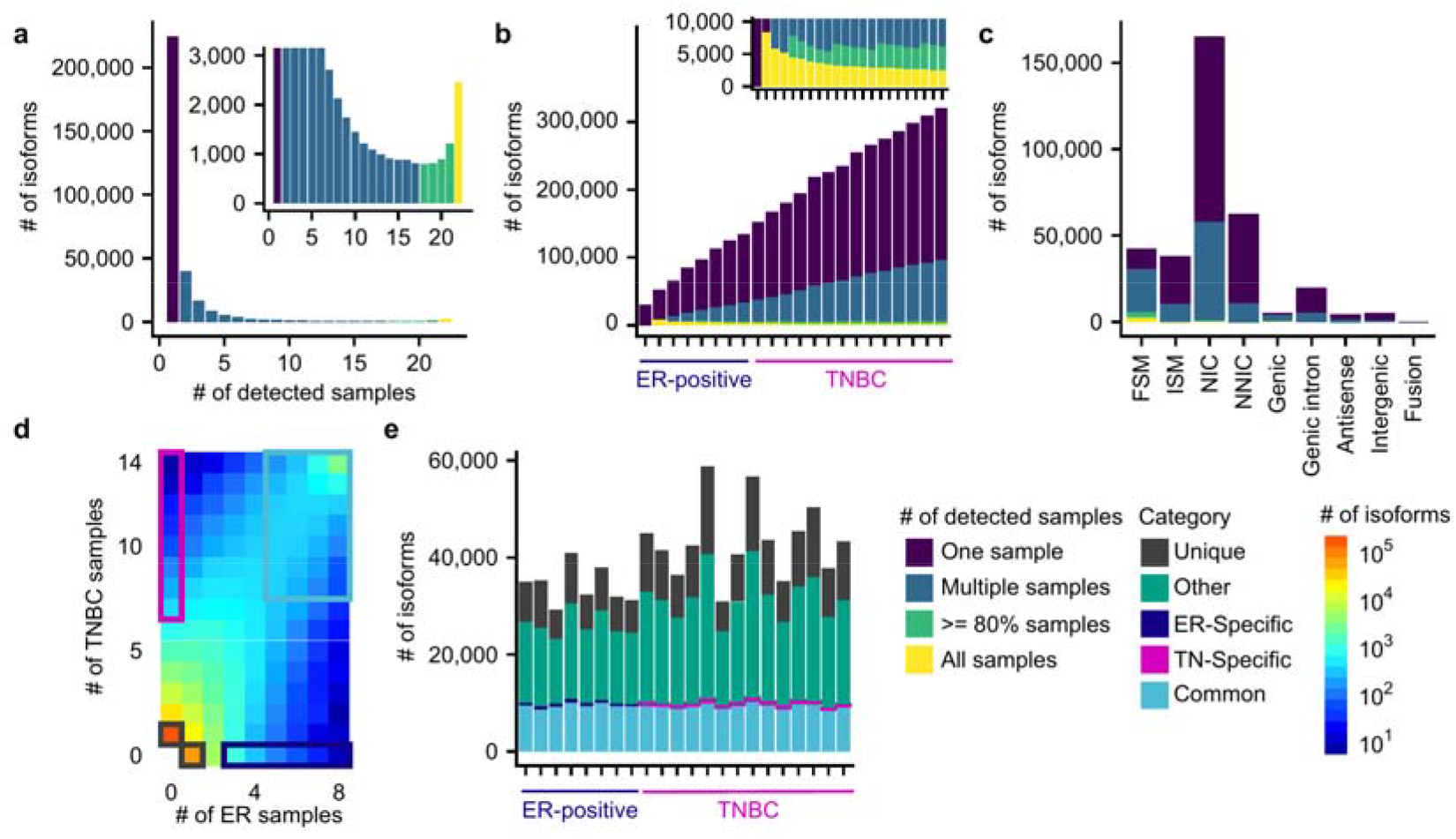
Isoform distribution detected by MuSTA. **a**, The number of isoforms according to the number of samples that generated isoforms. **b**, The number of isoforms when incrementing sample numbers one by one. **c**, The number of isoforms for each SQANTI category which represents the similarity against reference transcripts. FSM, full-splice match; ISM, incomplete-splice match; NIC, novel-in-catalog; NNIC, novel not in catalog. **d**, A heatmap representing the distribution of the number of isoforms from ER-positive and triple negative subtypes. The group of isoforms indicated by light blue, blue, and black rectangles were defined as common, specific, and unique, respectively. **e**, The distribution of isoforms according to their categories defined in **d**.

Next, we further investigated this heterogeneity with the aid of SQANTI, which classifies isoforms into nine categories by comparing with reference gene annotation; FSM, incomplete-splice match (ISM), NIC, novel not in catalog (NNIC), genic, genic intron, antisense, intergenic, and fusion. FSM and ISM only contained known splicing junctions. NNIC consisted of isoforms with novel splicing donors or acceptors. We found NIC to have the highest number, and the pairing of splicing donors and acceptors found to be much more diverse than what we could find in the GENCODE. The second most found category was NNIC. While 80% of those NNIC were detected only in one sample, there were a certain number of isoforms recurrently detected. 2,765 isoforms were found in all samples, and most of them were classified as FSM. To the contrary, almost all isoforms that were classified as genic intron, antisense, intergenic, and fusion were detected only in one sample (Fig. 3c). Fig. 3d shows the number of detected isoforms according to the number of samples of each subtype. Isoform detection itself, as well as isoform expression, was shown to be informative for differentiating the two subtypes (Supplementary Fig. 8). Therefore, we classified the isoforms into four categories; isoforms found in more than half of the samples in both subtypes were defined as “common”; isoforms found in only one subtype and the sample number being significant (p<0.05) in two-tailed Fisher’s exact test (i.e., more than three samples in ER-positive breast cancer, and more than seven samples in TNBC) were defined as “specific” (Supplementary Table 2); isoforms detected in only one sample were defined as “unique”; the rest of isoforms were defined as “other”. Whereas the number of unique isoforms varied according to the total number of isoforms in each sample, there was little variation in the number of common isoforms. Specific isoforms were in the range of 100-200 isoforms per sample (Fig. 3e).

### Subtype-specific isoforms

We hypothesized that subtype-specific isoforms may contain key molecules for cellular pathways activated specifically in the corresponding subtypes. To address this, we selected top 100 subtype-specific isoforms with the highest transcript per million (TPM) fold change (Fig. 4a). Isoforms from key oncogenes in the ER-positive subtype, such as an *ESR1* isoform and a *PGR* isoform were present in the top 100 isoforms. The *ESR1* isoform is annotated in GEOCODE and the *PGR* isoform was a novel isoform. Also, in the figure were NIC isoforms from subtype-specific genes, including an *AGR3* isoform in ER-positive breast cancer^38^ and a *GABRP* isoform in TNBC^39^. As of NNIC isoforms, *KLK5* and *LOXL4* have been reported to be a tumor suppressor gene (TSG)^40^ and be related to the metastasis in breast cancer^41^ respectively, and four isoforms of these genes were among the top 100 subtype-specific isoforms as well. It is likely that the number of subtype-specific isoforms reflected the association of the genes with the respective subtypes. In reality, *ESR1* had the largest number of subtype-specific isoforms among ER-positive breast cancer (Fig. 4b). *GABRP* had the largest number of subtype-specific isoforms in TNBC, and other oncogenes such as *BCL11A* and *PABPC1* also had many TNBC-specific isoforms. The results of this analysis may lead to the identification of novel oncogenes associated with breast cancer. For example, seventeen isoforms of unknown origin (“novel genes”) were of the top 100 isoforms, that warrant further investigation.

**Figure 4.**
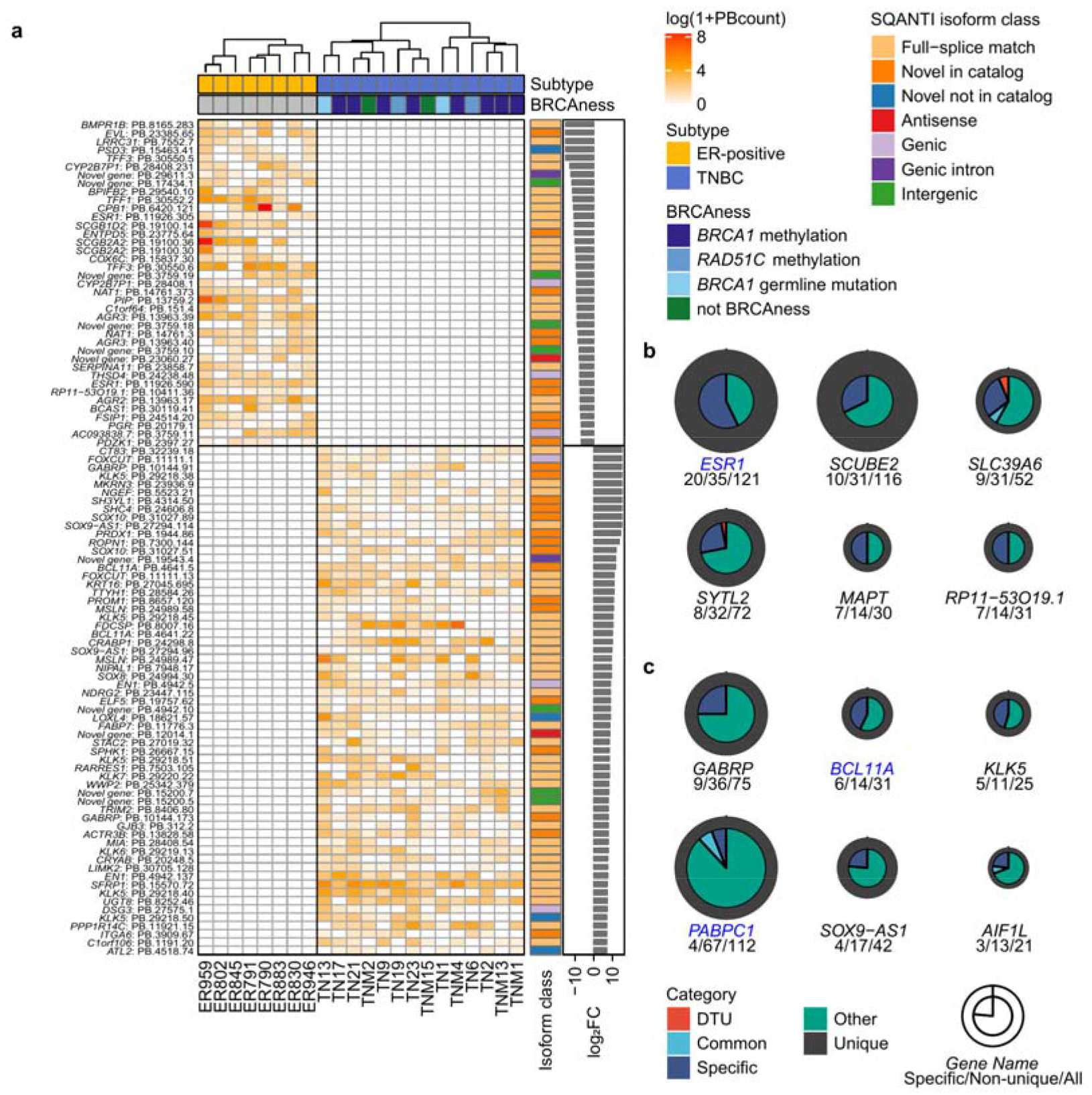
Subtype-specific isoforms. **a**, Top 100 subtype-specific isoforms with the highest fold change of transcript per million (TPM). Log-transformed PBcount data in each sample are indicated in a heatmap. Subtype and BRCAness are annotated for samples (top), and isoform classification by SQANTI and TPM fold change are annotated for isoforms (right). Absolute values of fold change larger than 10 are truncated. **b**, Circular plots indicating the number of isoforms according to isoform categories defined in Fig. 2D and differential transcript usage (DTU). Gene symbols are colored in blue if genes are oncogenes. Three numbers below each gene symbol mean the number of specific (left) / non-unique (center) / all (right) isoforms, respectively.

In order to investigate the true existence of isoforms detected in MuSTA, we focused on *SOX9-AS1*. *SOX9-AS1* is a long non-coding RNA that exists in the antisense of SOX9, a transcription factor that regulates cell differentiation for organ development in the fetal stage^42^. In our data, two isoforms from *SOX9-AS1* were expressed strongly in TNBC (Fig. 4a), and forty-two isoforms including four TNBC-specific isoforms were detected (Fig. 4b). We detected isoforms with read-through transcripts from *SOX9-AS1* and *AC005152.3*, which is located adjacent to *SOX9-AS1*. Using nested PCR, we were able to confirm the existence of these isoforms (Supplementary Fig. 9).

### Differential transcript usage in MuSTA-transcriptome

As another approach to capture subtype-related isoforms, we conducted DTU tests with the transcriptome obtained by MuSTA, under the assumption that those genes have functional relevance to the breast cancer biology (Fig 5, Supplementary Fig. 10).

**Figure 5.**
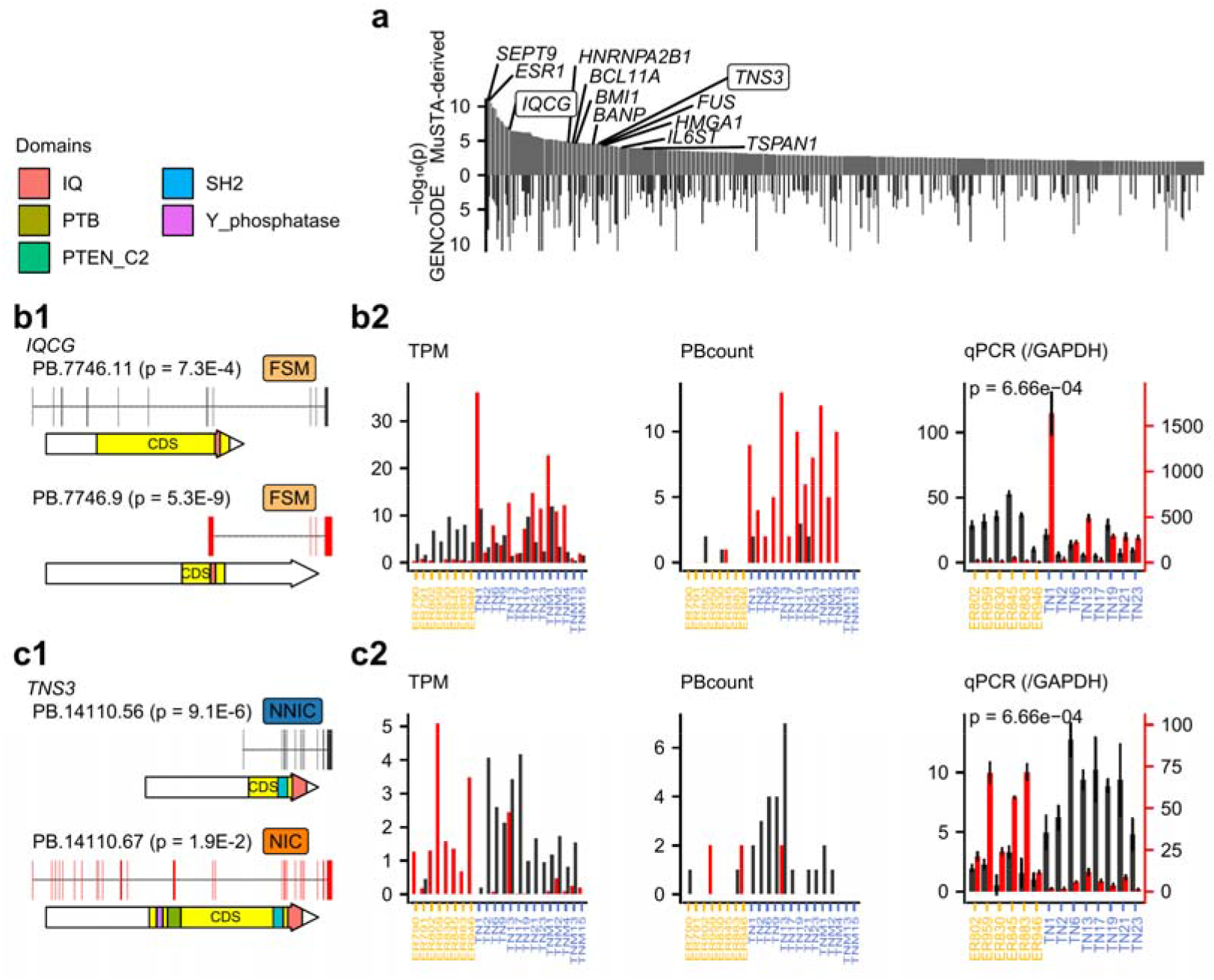
Differential transcript usage in MuSTA transcriptome. **a**, of differential transcript usage (DTU) inference in isoforms with p < 0.01 when using MuSTA-transcriptome. P-values were corrected in a stage-wise manner described in ref. 61. for GENCODE annotation are shown if isoforms are annotated in GENCODE. We sorted genes according to p-values for the probability of DTU in ascending number based on MuSTA-transcriptome. Ten gene symbols with the smallest p-values are labeled if genes are reported to be oncogenes or TSGs. Two genes were validated by qRT-PCR and are labeled with boxes. **b** and **c**, qRT-PCR validation of *IQCG* (**b**) and *TNS3* (**c**). Left, SQANTI classifications, transcript structures, and predicted protein domains of two DTU isoforms with the smallest p-values. Right, expression of DTU isoforms. Three types of expression data are shown (i.e. transcript per million (TPM) aligned to the MuSTA-derived transcriptome, PBcount, and relative qPCR expression against *GAPDH*). Relative qPCR expression has two y-axes along with DTU isoforms, because qPCR was conducted separately for each isoform. Error bars in qPCR expression indicates standard deviation of three replication studies. P-values were calculated with two-tailed Mann-Whitney U test for relative expression of DTU isoforms.

In order to examine the functional significance of DGE and DTU genes, we conducted gene ontology and KEGG pathway enrichment analysis (Supplementary Fig. 10d, e). In DGE genes, the following pathways and processes were enriched; biological process of peptidase and endopeptidase regulation, molecular function associated with ectodermal development and differentiation, and KEGG pathway associated with cell cycle. Curiously, eight out of the top ten biological processes enriched in DTU genes were associated with molecular binding. mRNA metabolic process, cell division, and RNA processing were included in the molecular functions enriched in DTU genes. Spliceosome and Cell cycle were the two significantly enriched KEGG pathways. Therefore, isoform switching was suggested to regulate cell cycle and RNA management. Of note, while the DTU genes and particularly the DTU isoforms were not identical between MuSTA-transcriptome and GENCODE-transcriptome, these findings remained the case (Supplementary Notes, Supplementary Fig. 11).

RT-qPCR confirmed that DTU was indeed observed for *IQCG* and *TNS3* (Figs. 5b and c), both of which have reported to be associated with breast cancer or other malignancies^43–47^. While the DTU isoforms of *IQCG* were matched to a previous report^43^, unannotated isoforms were included in the DTU isoforms of *TNS3*. The NNIC isoform of *TNS3* (PB.14110.56) had an unannotated first exon, and the genomic sequence of this exon was conserved among vertebrates (Supplementary Fig. 12). In addition, this first exon was associated with the peaks of three chromatin modifications, H3K4me3, H3K27ac, and H3K4me1, in MDA-MB-468 and MCF-10A, cell lines of TNBC and normal breast epithelium, whereas these peaks were not observed in MCF-7, an ER-positive cell line. The former modification is known to be enriched at promoters, and the latter two are used as enhancer markers. These findings reinforce the existence of the unannotated isoform and suggest that it was under the control of epigenetic regulation.

Thus, through the detection of DTU genes, we could successfully identify the genes with implication in breast cancer biology. Therefore, it may be possible to identify genes that play important role in breast cancer through further analysis of the DTU genes detected with MuSTA.

### Transcript structure of fusion transcripts

While short-read sequencing makes it possible to detect the break points of structural variation with high sensitivity and accuracy, long-read sequencing enables us to see the structure of resultant transcripts accurately to the extent that could not be achieved with short-read sequencing.

Of the chimeric IsoSeq cluster reads found in nine TNBC samples, we identified 402 reads with corresponding break points in whole genome sequencing (WGS) data (Supplementary Fig. 13). When the transcript fragments 5’ and 3’ of the fusion points were multi-exonic, almost all were mapped to genic region. In contrast, when they were mono-exonic, more than half were mapped onto non-genic regions (intergenic, genic intron, or antisense regions) (Figs. 6a and 6b). Almost all transcript fragments with TSS were mapped to genic regions, while only half of the downstream fragments were mapped to genic regions, possibly reflecting that the initiation of transcription needs known TSS with stringent requirement.

**Figure 6.**
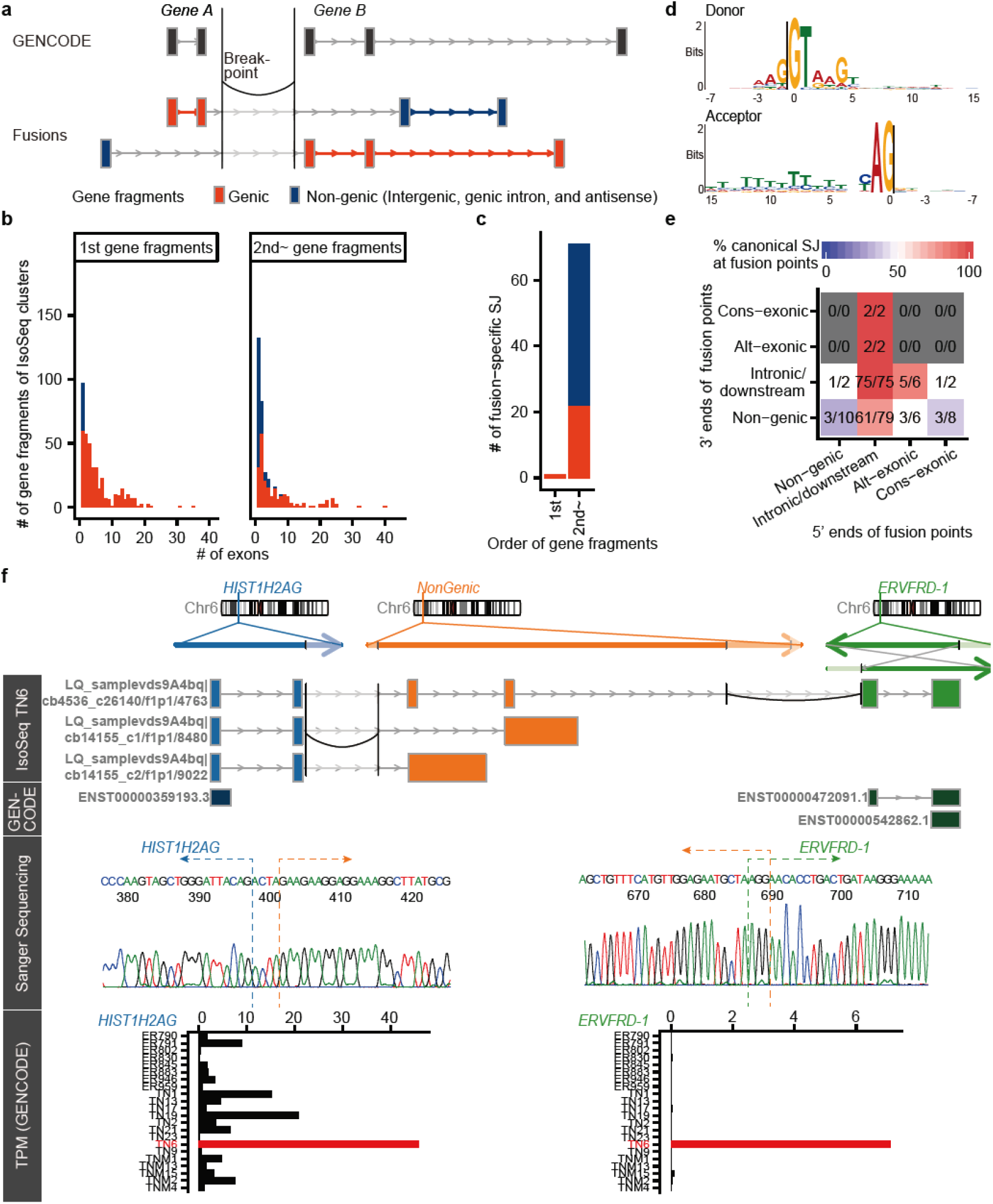
Transcript structure of fusion transcripts. **a**, A schematic of gene fragments. **b**, Distribution of the number of exons per gene fragments according to their genomic regions and their order in fusion transcripts. **c**, Bar plots showing the number of fusion-specific splicing junctions for every order of gene fragments in fusion transcripts. **d**, The motif of fusion-specific splicing junctions. We note that the first intronic two bases were intentionally chosen. **e**, A heatmap with axis representing exon-intron structures at both edges of fusion points. Colors and numbers indicate the proportion of confirmed canonical splice sites at fusion points. Non-genic means that the corresponding gene fragments were mapped to intergenic, genic intron and antisense regions. Intronic/downstream means a fusion point is on an intron or downstream of genes. Alt-exonic means alternative exon, and cons-exonic means constitutive exon. **f**, Structure of three-piece fusion transcripts from *HIST1H2AG-NonGenic-ERVFRD-1*. The genomic axes represent three original genomic regions. Below them are chimeric IsoSeq cluster reads. Curves correspond to structural variants detected with whole genome sequencing data. The category “GENCODE” shows annotated transcripts. Outside regions of structural variants are shaded. Exon-intron structures don’t necessarily reflect accurate length for visibility. TPM denotes transcript per million.

Next, in order to characterize the aberrant transcription caused by chromosomal rearrangement, we examined fusion-specific splicing junctions; we’ve defined fusion-specific splicing junctions as splicing junctions that exist on neither GENCODE transcripts nor non-chimeric MuSTA isoforms. Most of them were located in 3’ of the fusion points (Fig. 6c). 49 (68%) of fusion-specific splicing junctions were found from non-genic regions, while some were also detected from genic regions. The motif of fusion-specific splicing junctions was similar to that of ordinary canonical junctions (Fig. 6d), while the caveat is that the first intronic two bases were intentionally chosen because we removed the chimeric reads with the non-canonical junctions (other than GT-AG, GC-AG, and AT-AC) which were not detected in GENCODE or MuSTA-transcriptome. When we sorted the reads according to genomic regions of fusion points (Fig. 6e), the 5’ ends of fusion points mostly existed within introns or downstream of the genes to which the fusion transcript fragments were assigned. When both sides of fusion points were located at introns or downstream of genes, they agreed with canonical splicing motif in all cases (75/75), indicating that novel exons or splice junctions concordant with the splicing rules were indeed generated in association with chromosomal rearrangements. Although only few were found, there were reads that agreed with splicing motif even when 5’ ends or 3’ ends of fusion points existed on constitutive exons. It can be speculated that exon-intron structures have changed by structural context in these reads.

### Three-piece Fusion transcripts with complex structural variations

In recent years, long-read genomic sequencing has been suggested to identify complex structural variations (SV) which was impossible to detect with next generation sequencing^48,49^. However, the question of what kind of transcript can be found from complex SV is yet to be known. Of the chimeric IsoSeq reads, we’ve identified five non-redundant reads that were both mapped to three regions and had break points that were detected in WGS data (Fig. 6f, Supplementary Fig. 14, Supplementary Table 4). We’ve confirmed the fusion transcripts of *HIST1H2AG-NonGenic-ERVFRD-1*, *OGG1-NonGenic-NonGenic*, and *SLC12A2-NonGenic-SLC12A2* by sanger sequencing of PCR amplicon. It is worth noting that two fusion reads, *HIST1H2AG-NonGenic-ERVFRD-1* and *SMIM13-NonGenic-NonGenic* were transcribed from the sense and antisense strands of the same rearranged locus, respectively, though we could not amplify the latter fusion transcript by PCR. Very recently, the Pan-Cancer Analysis of Whole Genomes consortium has found several bridged fusion transcripts which were mapped to two genomic regions that were connected by untranscribed intervening genomic fragment^2^. However, the three-piece fusions we found had internal genomic regions of more than thousands of base pairs and some fusions were even spliced in these regions. This type of fusion transcripts can’t be found without long-read transcriptome sequencing.

Because *HIST1H2AG-NonGenic-ERVFRD-1* contains full CDS of *ERVFRD-1*, full length ERVFRD-1 protein might be translated from the fusion transcript. *ERVFRD-1* was specifically expressed in the sample carrying the fusion transcript. Expression of *ERVFRD-1* is generally suppressed across all tissues except for placental tissues^50^. Further, considering the chromatin modification status, it is quite possible that this three-piece fusion transcript utilized the cis-regulatory region of *HIST1H2AG* observed in one normal breast epithelial cell line and two breast cancer cell lines (Supplementary Fig. 15), suggesting the existence of a mechanism like enhancer hijacking^51^. The largest difference from enhancer hijacking is that the promoter and enhancer region of *HIST1H2AG* was located 50kb upstream of *ERVFRD-1* even in the rearranged chromosome, and *HIST1H2AG-NonGenic-ERVFRD-1* exploited them by forming the readthrough transcript that strided over two break points.

We also noticed that *ERVFRD-1* was highly expressed in 4 samples from TCGA that carry *ERVFRD-1* fusions (*ABHD12-ERVFRD-1*, *ELOVL2-ERVFRD-1*, *NEDD9-ERVFRD-1*, and *NOL7-ERVFRD-1*, respectively, according to FusionGBD^52^), and amplified in several types of cancers, including melanoma, uterine cancer, and ovarian cancer (Supplementary Fig. 16). Because it was reported that *ERVFRD-1* suppressed the antitumor immunity in a syngenic mouse tumor model^53^, the aberrant expression of *ERVFRD-1* we observed might contribute to the oncogenesis.

## Discussion

Although next-generation sequence technology allows comprehensive survey of RNA alterations, it was possible merely to focus on individual splicing junctions depended on existing reference annotation. Previously, it was difficult to acquire non-redundant annotation from long-read RNA sequencing or apply them to multiple samples, preventing researchers from conducting inter-subtype comparison such as DGE and DTU. We have here introduced MuSTA, a unique method for constructing full length transcriptome from multiple samples, and we showed through simulations that it enables more accurate evaluation of transcript usage. By utilizing MuSTA, we obtained 344,504 non-redundant isoforms including 263,711 unannotated ones from 22 breast cancer clinical specimens, unveiling the diversity and the heterogeneity of transcripts in both inter-subtype and intra-subtype levels. Subtype-specific isoforms with high TPM fold change contained unannotated isoforms from subtype-specific biomarkers and novel genes, which may play important roles in the carcinogenesis of each subtype. The transcriptome determined with MuSTA allowed us to conduct comprehensive DTU analysis using isoforms expressed in target sample groups. We identified DTU of *IQCG* and *TNS3* and validated them using qRT-PCR.

Bjørklund SS *et al.* have reported that *IQCG* intronic TSS induces exons 9-12 overexpression in ER-negative breast cancer^43^. They pointed out that because this region can be translocated and fused to the N-terminal of *NUP98* in an acute T-lymphoid/myeloid leukemia^44^, deregulated expression in this region may be oncogenic. We’ve confirmed that PB.7746.9 (matched to ENST00000478903.5) was overrepresented compared to other isoforms in TNBC, and this indeed matched the intronic-start transcript they’ve reported (Fig 4b).

*TNS3* (Tensin3) is a protein with SH2 domain and C2 domain, and is known to contribute to cell migration, anchorage-independent growth, and metastasis in several types of cancer including breast cancer^45–47^. Quan X *et al.* reported that tumorigenesis and metastasis are introduced by Src-induced phosphorylation of its SH2 domain^45^, and it has been reported that epidermal growth factor (EGF) or platelet-derived growth factor (PDGF) provokes phosphorylation of C2 domain in TNS3, resulting in the activation of Rho GTPases to drive directional cell migration^46^. EGF is also shown to suppress TNS3 and upregulate C-terminal Tensin-like protein (CTEN) simultaneously and to promote mammary cell migration^47^. The isoform prominent in TNBC (PB.14110.56) lacked N-terminal C2 domain and have similar structure with CTEN, implying that the isoform may regulate the function of full length TNS3. By combining the result with recent studies, we concluded that isoform switching might have oncogenic effects in both genes.

Furthermore, we exploited the full-length sequencing capability of IsoSeq to show several features of exon-intron structures of fusion transcripts and fusion-specific splice sites. We also detected three-piece fusion transcripts that were transcribed from three distinct genomic regions.

To summarize, full-length transcript sequencing in multiple samples provides a complementary transcript-level analyses on top of the conventional RNAseq approaches, enabling us to focus on the isoforms from target cells, and apply pre-existing analyses such as clustering, DGE, DTU, and gene ontology analysis. MuSTA requires reference genomic sequence and transcriptome data as the mandatory input other than long-read RNA sequencing data, and these are available for human and other several species. There are two long-read sequencing techniques, SMRT sequencing from Pacific Biosciences and Oxford Nanopore Technologies (ONT) MinION. Although we used SMRT sequencing reads in this paper, MuSTA can be applied to ONT reads in theory.

Our methods have the potential for a variety of utilizations. Spliceosome disruption sometimes plays key oncogenic roles such as recurrent mutations on U1 snRNA in medulloblastoma and other cancers^54,55^. By applying MuSTA to cancer types with cryptic splicing, we will also be able to look into transcriptome wide changes. Since there is emerging evidence that gene isoforms are responsible in cancer for survival^56^ and drug response^57^, elucidating cancer profiles at isoform level might provide further druggable targets or previously undiscovered biomarkers. In recent years, cancer-specific splicing junctions are investigated comprehensively, and are considered as an important source of neoantigens^58^. By applying MuSTA in a large scale, it might be possible to look into how these junctions bring impact in transcript level, and that might lead to the development of immunotherapies. Even in diseases other than cancer, there are increasing evidence that variant-induced splice alteration might lead to diseases such as coronary artery disease and multiple sclerosis etc…. It is said that sQTL interferes with the phenotypic traits in at least the same level, if not more, as expression QTL^59^. Focusing on transcript level might enable us to find the relationship with variants and disease that haven’t been identified before.

There are several limitations in our analysis. First, the biggest limitation is the lack of sample numbers to detect enough isoforms with low expression. Also, because it is difficult to efficiently amplify long (>6 kbp) transcripts, reading the whole length of long transcript might result in insufficient processing, meaning only part of the transcript might be read. Currently the technologies of long-read sequencer are developing rapidly. For example, a newly developed “Sequel II” generates at least 8 times more data and far longer raw reads than the conventional Sequel, according to Pacific Biosciences (https://www.pacb.com/blog/award-winning-sequel-ii-system). The advance of SMRT sequencing will improve the comprehensiveness of isoform detection in MuSTA, and the analysis of transcript level expression can be expected to increase in its accuracy. Second, although MuSTA provides cohort-wide transcriptome which contains a large number of unannotated isoforms, the procedure was not fully annotation-free, as MuSTA employs SQANTI filtering, which compares the transcriptome with a reference gene annotation to exclude potential artifacts. Third, whereas we mainly targeted the isoforms shared across several samples, each sample contained substantial number of unique isoforms, which deserves further evaluation. These isoforms may reflect sample specific states including somatic mutation and/or epigenetic alteration^60^, or may merely be the resultant of aberrant splicing coupled with elevation of gene expression. The fourth limitation is the difficulty of examining TSG. We’ve identified many subtype-specific oncogenes, potential oncogenes, and biomarkers, but that was not the case for TSG. Many TSGs were low in expression, and it was difficult to conduct accurate quantification through RNAseq; even isoform detection became challenging in long-read sequencing technique. Therefore, even by using the method here, it was difficult to conduct transcript level analysis for TSG. Finally, as we established the strategy to obtain cohort-wide transcriptome by focusing on the consistency of individual splice sites, differences in TSS or transcript termination sites (TTS) were not fully investigated in this paper, although this information was retained throughout the MuSTA process; there might be more suitable strategies for summarizing transcripts from multiple samples when surveying TSS and TTS.

Despite these limitations, our results demonstrate that MuSTA is a useful tool to complement future transcriptome studies. Our findings are impossible to find through other methods, and this method, MuSTA, can contribute to the further understanding of cancer biology and many other cell biology in transcriptome level.

## Materials and Methods

### Reference genome and annotation files

We used hg38 as the reference genome, and GENCODE version 28 comprehensive gene annotation as the annotation file unless otherwise noted. We focused on isoforms mapped to autosome or chromosome X after we applied our pipeline.

### Definition of oncogenes and tumor suppressor genes (TSGs)

We defined oncogenes as those identified in at least one of three curated oncogene databases, cancer gene census (version 90), ONGene, and OncoKB (version 1.23). Since ONGene collected only oncogenes, we used the other two repositories for TSGs.

### Whole genome sequencing (WGS)

Whole genome sequencing (WGS) were conducted and reported in the previous report^31^. The original data were publicly available (https://humandbs.biosciencedbc.jp/en/hum0094-v3#WGS). In the present study, we re-analyzed the data as follow. We detected mutations and structural variants (SV) in two ways and combined them together. First, as shown in our previous research^31^, we used in-house pipeline for analysis. Since we worked with hg19 for our pipeline, the results were transferred to hg38 using liftover tool. Next, in order to detect short structural variants, we used Genomon^62^, an analytic pipeline for next-generation sequencing data, which carries out mapping using STAR^63^, annotation, and further functions including detection of SV for DNA and intron retention for RNA.

### RNAseq

Transcriptome sequencing (RNAseq) were conducted and reported in the previous report^31^. The original data were publicly available (https://humandbs.biosciencedbc.jp/en/hum0094-v3#RNA-seq). RNAseq was performed with 100 bp paired-end reads using NEBNext Ultra Directional RNA Library Prep Kit for Illumina (New England BioLabs, Ipswich, MA, USA) according to the manufacturer’s protocol. In the present study, we re-analyzed the data as follow. RNAseq reads were mapped to hg38 reference assembly and expression data was calculated in the format of transcript par million (TPM) by 2 ways, quasi-mapping-based mode of Salmon^64^ and STAR-RSEM protocol^63,65^. Salmon index was used with ‘–keepDuplicates’ option, and Salmon quant was performed with ‘-l A –gcBias – seqBias –validateMappings’ options. RSEM was performed with following commands,

- rsem-prepare-reference –gtf –star
- rsem-calculate-expression –star –paired-end <short-1.fastq> <short-2.fastq>

We detected splice junctions and intron retention using Genomon, which ran STAR internally for mapping to reference genome.

For Annotation file, we used GENCODE version 28 or a set of transcripts retrieved from our SMRT analysis pipeline.

### BRCAness

We defined the triple negative breast cancers with defective homologous recombination (BRCAness) based on the profiles of structural variations, mutational signature, germline mutational status of *BRCA1*, expression of *BRCA1* and *RAD51C*, and promoter methylation of these *BRCA1* and *RAD51C* in the previous report^31^.

### SMRT sequencing

Long-read sequencing was performed with the Pacific Biosciences Single-Molecule Real-Time (SMRT) sequencing technology with SMRT cell chemistry (SMRTbell Template Prep Kit 1.0, Sequel Binding Kit 2.0, Sequel Sequencing Kit 2.0, all from Pacific Biosciences, Menlo Park, CA, USA). Full-length cDNA libraries were constructed from 1 μg of total RNA with SMARTer® cDNA synthesis kit (Takara Bio, Kusatsu, Shiga Japan), utilizing switching mechanism at 5’ end of RNA template (SMART) technology coupled with PCR amplification. PCR amplification was performed with PrimeSTAR® GXL DNA Polymerase (Takara Bio). The sequencing templates used for SMRT sequencing on the Sequel platform (SMRTbell) were constructed from 1 μg of PCR products. After DNA damage and end repair, the SMRTbell adaptors were ligated onto the PCR amplicons, followed by purification with 0.6 volumes of Agencourt AMPure PB (Pacific Biosciences) with a 10-minute incubation on a vortex mixer. Primer annealing and DNA polymerase binding were carried out according to the manufacturer’s instructions. Briefly, sequencing primers were annealed to the template at a final concentration of 0.833 nM by denaturing the primer at 80°C for 2 minutes and cooling to the 4°C before incubating with the library at 20°C for 30 minutes. Distributions of the SMRTbell size were presented in Fig. 5.

This library went through sequential DNA replication, with DNA polymerase detachment as replication limitation, and was analyzed by IsoSeq2 pipeline using SMRTlink^15^ with following settings; maximum dropped fraction was set to 0.8, maximum subread length of 15000, minimum subread length of 50, minimum number of passes of 1, minimum predicted accuracy of 0.8, minimum read score of 0.65, minimum SNR of 3.75, minimum Z score of −9999, minimum quiver of 0.99, trim QV’s 3’ of 30, and trim QV’s 5’ of 100, minimum sequence length of 200, polish CCS as false, emit individual QVs as false, and required polyA as true. In IsoSeq, consensus reads “Read of Insert (RoI)” were obtained. RoIs with both cDNA primers and poly(A) were defined as full-length (FL) reads, and others were defined as non-full-length reads. IsoSeq clustered these reads into isoform sequences using an algorithm called ICE. “Polished” reads from the algorithm (specifically, polished_hq.fastq and polished_lq.fastq as output file names of IsoSeq2) were put into further analyses. Of note, those reads do not necessarily represent non-redundant isoforms because of ICE characteristics and natural 5’ degradation in RNA.

### Hybrid error correction

We used LoRDEC^17^ for hybrid error correction of IsoSeq reads with RNAseq data. LoRDEC was executed with following commands,

- lordec-build-SR-graph -T 3 −2 <RNAseq_interleaved.fastq> -k 19 -s 3 -g
- lordec-correct -T 8 -i <Isoseq_reads.fastq> -k 19 -s 3 −2 <RNAseq_interleaved.fastq> -o <corrected_Isoseq.fastq>

### Mapping of corrected IsoSeq reads

Next, IsoSeq reads were mapped to hg38 reference assembly by Minimap2^66^ with similar two commands to retrieve results as both SAM and PAF formats under the same condition,

- minimap2 -ax splice -uf -C5 –secondary=no <GRCh38.mmi> <corrected_Isoseq.fastq>
- minimap2 -cx splice -uf -C5 –cs –secondary=no <GRCh38.mmi> <corrected_Isoseq.fastq>

We filtered IsoSeq reads with mapping quality larger than 50.

### Intra-/inter-sample collapsing IsoSeq reads

Intra-sample integration of mapped IsoSeq reads, followed by inter-sample integration, was performed by our R code. As for multi-exon transcripts, we merged IsoSeq reads with same splice junctions. The most upstream TSS and the most downstream TTS of original transcripts of merged isoforms were defined as the TSS and TTS of the merged isoforms, with all the original TSS and TTS information linked and retained. We combined 5’ truncated multi-exon isoforms with longer and compatible isoforms in intra-samples (meaning, we treated them as fragments of longer transcripts.) On the other hand, we considered those truncated transcripts as independent transcripts of longer transcripts from other samples unless they shared all splice junctions, in order to detect correct exon-intron structure from transcripts expressed in target cells. Regarding single-exon transcripts, we consolidated reads with other single-exon transcripts if genomic range of the former transcript was overlapped with the latter read. We didn’t combine single-exon transcripts with longer multi-exon transcripts. Both intra-sample and inter-sample integration were performed according to this procedure.

We considered that an isoform was detected in a particular sample only when there were non-truncated IsoSeq reads in the sample (i.e. multi-exon IsoSeq reads of which all splice junctions matched, or mono-exon IsoSeq reads with genomic range within the integrated isoform). Isoform count was defined as the sum of IsoSeq reads FL count which were linked to a isoform and not to any other isoforms, and we named it as PBcount.

### Classification and Filtering with SQANTI

We have classified and filtered curated isoforms with SQANTI^67^. For classification, we used genomic range of isoforms in GTF format, TPM of RNAseq yielded by Salmon, and the number of FL reads summed in the last section. SQANTI uses random forest to identify whether an isoform is an artifact. As in the primary setting, isoforms with all splicing junctions matching those in annotated transcripts (full-splice match), were set as true positives in the training data; true negatives were defined as those transcripts with at least one novel and non-canonical splicing junction. We used isoforms that passed SQANTI filter as our full-length transcript library for downstream analyses.

### Chimeric reads

While chimeric reads are mapped onto more than two genomic regions, because long-reads yield certain amount of sequencing error, there might be some uncertainty of fusion sites within the range of several bp. Therefore, when we found only one position with canonical splice junctions, as long as it was within the uncertainty range of the fusion sites, we picked up the position as fusion points. For the reads with splice junctions as fusion points, we made an additional requirement for them to have genomic break points within 100,000bp of fusion points. As of the reads that weren’t confirmed, assuming that they had break points on exonic regions, we made a requirement that there are genomic break points within 100bp of fusion points. Of the three-piece fusion transcripts with two fusion points, we picked up those with one linked and the other one unlinked to a genomic break point. Following that, we manually searched for possible genomic break points that correspond with the unlinked fusion points with blastn^68^.

### Simlady, a simulator for long-read RNA sequencing

Contrary to long-read genomic sequencing where reads are generated from the distribution of read length, reads are generated from template transcripts in long-read RNA sequencing. The length of generated reads could be different from the original template transcripts, and we have focused on RNA 5’ degradation and sequencing error as its main reason. RNA decay is considered as a major reason why transcript start sites (TSS) of IsoSeq reads can be inaccurate^69^. As of transcript termination sites, it has been reported that there’s only a few amounts of error^70^, therefore, we didn’t investigate further. RNA decay was fitted by gamma distribution. The ‘pelgam’ function implumented in ‘lmom’ R package was used for fitting. A public data that uses MCF-7 cell lines for IsoSeq (https://github.com/PacificBiosciences/DevNet/wiki/IsoSeq-Human-MCF7-Transcriptome) is often used in simulator evaluation^28^; therefore, we’ve used this for evaluation. Universal Human Reference (brain, liver, and heart) IsoSeq data (https://github.com/PacificBiosciences/DevNet/wiki/Sequel-II-System-Data-Release:-Universal-Human-Reference-(UHR)-Iso-Seq), which is another public data, was used as a validation dataset. We looked into how much TSS of FLNC reads are shortened according to the nearest upstream TSS in GENCODE Using gamma distribution, the shortened length matched well in the range of <10,000bp (Supplementary Fig. 1). This distribution showed to be extremely heavy tailed, and hardly matched with fitted curve for more than 10,000bp. This may be because since GENCODE TSS annotation was imperfect, there were FLNC reads that incorrectly linked to distant reference TSS. As of sequence error model, we employed SimLoRD^28^ model. While SimLoRD inserts error in order not to change the read length, the read length changes actively according to the error inserted. Since SMRT sequencing data shows a transcript-length dependent distribution, each read is sampled according to the probability derived from fold change and transcript length, in order to re-present this distribution.

### Simulations with different settings

We generated two groups of short-read and long-read RNAseq data from the GENCODE annotation. We set the number of samples per groups as 8, fold change for DTU as 4, short-read length as 100bp, short-read depth as 50,000,000 reads, and the number of FLNC reads as 250,000. We changed these values one by one for investigating the effect on DTU inference (Supplementary Fig. 3). In detail, we assigned 1 as relative expression of all isoforms except for randomly selected 10% genes, for which we randomly selected 2 isoforms for DTU and assigned the pre-defined fold change value to one isoform in the first group and the other isoform in the second group. ‘Simulate_experiment_countmat’ function in polyester r package^71^ was employed for simulating short-read RNAseq data, and subsequently the reads were shuffled because the reads were written out by each transcript. We used simlady for simulating FLNC reads without specifying read length distribution, and simlady generated reads under the log-normal distribution inherited from SimLoRD^28^. The FLNC reads were then processed to cluster reads by ‘isoseq3 cluster’ with ‘–singletons’ option. While IsoSeq3 discards singletons, we determined the number of FLNC read suitable for IsoSeq2, which additionally uses non-full-length reads and tolerates clusters with only one FLNC read. Therefore, we combined singletons with clustered reads, and used them as input for MuSTA.

### Simulations based on the breast cancer dataset

We generated two groups of short-read and long-read RNAseq data from the FSM and NIC isoforms in the MuSTA-derived transcriptome obtained from 22 breast cancer specimens. The number of samples per groups was 8 and 14, respectively, which was the same number as the original data. We permutated the log-averaged expression of FSM isoforms and NIC isoforms separately. We assigned randomly DGE for 25% of all genes, and DTU for 10% of all genes so that 4% of genes were assigned as both DGE and DTU. These values were approximately the same as the original breast cancer data at an FDR threshold of 0.05. The expression fold change between groups was set to 4 for all isoforms of DGE genes so that the log-averaged expression remained the same. As for DTU genes, we shuffled all genes and tried to select two DTU isoforms, where one isoform had larger expression in the first group, and the other had larger expression in the second group. That is, we selected two isoforms with the highest expression in three ways at random, (i) two FSM isoforms, (ii) one FSM isoform and one NIC isoform, or (iii) two NIC isoform, so that NIC rate against all DTU isoforms reached the pre-defined value. Again, we set the expression fold change between groups to 4 for DTU isoforms so that the log-averaged expression remained the same. Short-read and long-read RNAseq reads were simulated as described above, with the exception that the length distributions of polished reads in breast cancer data were permutated and used for the FLNC read length distribution.

### Differential gene expression

Differential gene expression was investigated with DESeq2^72^ as described in ref. 73. Isoform expression data obtained with Salmon were imported into R and summarized at gene level using tximport^74^.

### Differential transcript usage

As Soneson C *et al.* compared state-of-the-art methods^11^, DEXSeq^75^ has shown to be the most accurate method, therefore, we chose DEXSeq as the inference engine of differential transcript usage in the way described by ref. 73, where in brief each isoform was treated as an exon, and a log-likelihood ratio test was performed under the setting with “~ sample + exon + subtype * exon” as the full model and “~ sample + exon” as the null model. For combining short-read RNAseq data and full-length PBcount data, we concatenated both data and set “~ sample + exon + subtype * exon + data type * exon” as the full model and “~ sample + exon + data type * exon” as the null model. We note the caveat that this setting treated RNAseq data and PBcount data derived from the same sample as biological replication, while DEXSeq does not have proper method for combining two technically replicated data with large batch effects, and we observed substantial difference between RNAseq data and PBcount data (Fig. 4). Though this could lead to artificial increase in power, on the contrary we got more conservative results from concatenated data than RNAseq data in simulations (Fig. 2, Supplementary Fig. 2). Gene-level and transcript-level false discovery rates were calculated with stageR^61^.

### Prefiltering of transcriptome

For pre-alignment prefiltering, we only retained those isoforms which had the first or second largest number of PBcount per gene in at least one sample and had equal with or more than five PBcount in all samples. Among these isoforms, up to 10 isoforms were chosen in descending order of PBcount. To avoid mismapping of RNAseq, for each gene with no selected isoforms, we also retained one isoform with the largest PBcount. We defined the selected isoforms as “major” isoforms. For post-alignment prefiltering, we used DRIMSeq^76^ filter and removed transcripts if they had proportion of expression lower than 0.1 compared to the total expression of the related genes.

### Overlap between MuSTA-transcriptome and unannotated open reading frames

The list of high-confidence translated open reading frames identified in human induced pluripotent stem cells and human foreskin fibroblast was obtained from ref. 37. We lifted the positions from Hg19 to Hg38, and retained those uniquely lifted. We counted the number of open reading frames whose genomic ranges were not overlapped with any GENCODE genes and were completely covered by MuSTA-transcriptome.

### Alternative splicing

We explored alternative splicing events of exon skipping/inclusion, alternative 5’, alternative 3’, mutually exclusive exons, and intron retention with ‘generateEvents’ function of SUPPA2^77^. Next, we used ‘performPCA’ function implemented in psichomics^78^ for principle component analysis of splicing events as described in a vignettes of the software (https://bioconductor.org/packages/release/bioc/vignettes/psichomics/inst/doc/CLI_tutorial.html).

### Domain prediction

We used HMMER^79^ against Pfam^80^ (version 32.0) with a following command, -hmmscan –domtblout –noali -E 0.1 –domE 0.01 Pfam-A.hmm

### Quantitative PCR and sequencing of fusion points

RNA was extracted from cells using an RNeasy Mini kit (Qiagen). Total cellular RNA was converted into cDNA by reverse transcription (SuperScript IV VILO Master Mix; Thermo Fisher) using random primers. Quantitative real-time PCR was performed using Power SYBR Green qPCR SuperMix-UDG with ROX (Thermo Fisher) through 40 cycles of 95°C for 15 sec and 60°C for 60 sec using an Applied Biosystems PRISM 7900 Sequence Detection System. Complementary DNAs for fusion points were amplified by reverse transcription PCR (RT-PCR) from RNA samples and subjected to Sanger sequencing. Primer sequences are provided in Supplementary Table 5.

### Chromatin modifications in breast normal/cancer cell lines

ChIP-seq of chromatin modifications for MCF-7, MDA-MB-468, and MCF-10A were carried out by Franco HL *et al.*^60^, and subsequently collected, mapped to hg19, and peak called by ChIP-Atlas^81^. We used the mapped data and the peak data with the threshold of q < 10^-5, and lifted them to hg38.

## Code availability

The source codes for our analyses are available upon request to the authors.

## Software version

We used the bioinformatic tools listed below: Genomon^62^ (version 2.6.0), STAR^63^ (version 2.5.2a), RSEM^65^ (version 1.3.1), Salmon^64^ (version 0.12.0), SMRTlink^15^ (version 5.1.0.26412), LoRDEC^17^ (version 0.9), Minimap2^66^ (version 2.12-r847-dirty), SQANTI^67^ (version 1.2), DRIMSeq^76^ (version 1.10.1), DESeq2^72^ (version 1.22.2), DEXSeq^75^ (version 1.28.3), tximport^74^ (version 1.10.1), stageR^61^ (version 1.4.0), HMMER^79^ (version 3.1b2), SUPPA2^77^ (version 2.3), psichomics^78^ (version 1.8.2), and BLAST^68^ (version 2.9.0+).

## URLs

COSMIC, https://cancer.sanger.ac.uk/cosmic; OncoKB, https://www.oncokb.org/; ONGene, http://ongene.bioinfo-minzhao.org/; cBioPortal, https://www.cbioportal.org/; GENCODE, https://www.gencodegenes.org/; Pfam, https://pfam.xfam.org/; HMMER, http://hmmer.org/; Pan-Cancer Analysis of Whole Genomes (PCAWG), https://dcc.icgc.org/pcawg; LoRDEC, http://www.atgc-montpellier.fr/lordec/; minimap2, https://github.com/lh3/minimap2; SQANTI, https://bitbucket.org/ConesaLab/sqanti/; SimLoRD, https://bitbucket.org/genomeinformatics/simlord/; R statistical software, https://cran.r-project.org/.

## Acknowledgements

We thank Ms. Miki Tamura, Ms. Kaori Sugaya, Dr. Manabu Soda, and Dr. Yoshihiro Yamashita for technical assistance. We are grateful to all the patients and families who contributed to this study. Computation time was provided by the Supercomputer System, Human Genome Center, the Institute of Medical Science, the University of Tokyo. This study was supported by the grants from Japan Agency for Medical Research and Development (AMED) (JP17am0001001 to H. M.; JP15cm0106085 to S.H.; JP19cm0106502 to M.K.,), the grant from the Japan Society for the Promotion of Science (JSPS) (16K07143 to M.K.), and the grant from the UBE Industries Foundation (to M.K.).

## Author contributions

S.N. developed the MuSTA pipeline and conducted experiments and data analysis. M.K. conducted experiments and supervised the study. T.O. and S.H. provided clinical specimens. Y.T., S.I., and F.K. prepared sequencing libraries and conducted experiment. T.U. and S.K. processed and analyzed sequenced data. S.N. and M.K. wrote the manuscript with comments from Y.S. and H.M.

## Competing interests

The authors declare no conflicts of interest.

## Notes

### Competing Interest Statement

The authors have declared no competing interest.

## References

1. Kim, J. & Eberwine, J. RNA: State memory and mediator of cellular phenotype. Trends in cell biology 20, 311–8 (2010).

2. Calabrese, C. et al. Genomic basis for rna alterations in cancer. Nature 578, 129–136 (2020).

3. Danan-Gotthold, M. et al. Identification of recurrent regulated alternative splicing events across human solid tumors. Nucleic Acids Research 43, 5130–5144 (2015).

4. Climente-González, H., Porta-Pardo, E., Godzik, A. & Eyras, E. The Functional Impact of Alternative Splicing in Cancer. Cell Reports 20, 2215–2226 (2017).

5. Biswas, K. et al. Intragenic DNA methylation and BORIS-mediated cancer-specific splicing contribute to the Warburg effect. Proceedings of the National Academy of Sciences 114, 11440–11445 (2017).

6. Grelet, S. et al. A regulated PNUTS mRNA to lncRNA splice switch mediates EMT and tumour progression. Nature Cell Biology 19, 1105–1115 (2017).

7. Salton, M. et al. Inhibition of vemurafenib-resistant melanoma by interference with pre-mRNA splicing. Nature Communications 6, 7103 (2015).

8. Chang, K. et al. The Cancer Genome Atlas Pan-Cancer analysis project. Nature Genetics 45, 1113–1120 (2013).

9. Shiraishi, Y. et al. A comprehensive characterization of cis-acting splicing-associated variants in human cancer. Genome research 28, 1111–1125 (2018).

10. Farver, C. et al. Comprehensive Analysis of Alternative Splicing Across Tumors from 8,705 Patients. Cancer Cell 34, 211–224.e6 (2018).

11. Soneson, C., Matthes, K. L., Nowicka, M., Law, C. W. & Robinson, M. D. Isoform prefiltering improves performance of count-based methods for analysis of differential transcript usage. Genome Biology 17, (2016).

12. Dueck, H. et al. Deep sequencing reveals cell-type-specific patterns of single-cell transcriptome variation. Genome biology 16, 122 (2015).

13. Tilgner, H., Grubert, F., Sharon, D. & Snyder, M. P. Defining a personal, allele-specific, and single-molecule long-read transcriptome. Proceedings of the National Academy of Sciences 111, 9869–9874 (2014).

14. Rhoads, A. & Au, K. F. PacBio Sequencing and Its Applications. Genomics, Proteomics and Bioinformatics 13, 278–289 (2015).

15. Gordon, S. P. et al. Widespread polycistronic transcripts in fungi revealed by single-molecule mRNA sequencing. PloS one 10, e0132628 (2015).

16. Koren, S. et al. Hybrid error correction and de novo assembly of single-molecule sequencing reads. Nature Biotechnology 30, 693–700 (2012).

17. Salmela, L. & Rivals, E. LoRDEC: accurate and efficient long read error correction. Bioinformatics 30, 3506–3514 (2014).

18. Abdel-Ghany, S. E. et al. A survey of the sorghum transcriptome using single-molecule long reads. Nature communications 7, 11706 (2016).

19. Wang, B. et al. Unveiling the complexity of the maize transcriptome by single-molecule long-read sequencing. Nature Communications 7, 1–13 (2016).

20. Sadler, K. C. et al. High resolution annotation of zebrafish transcriptome using long-read sequencing. Genome Research 28, 1415–1425 (2018).

21. Tilgner, H. et al. Comprehensive transcriptome analysis using synthetic long read sequencing reveals molecular co-association of distant splicing events. Nature biotechnology 33, 736–742 (2015).

22. Gupta, I. et al. Single-cell isoform RNA sequencing characterizes isoforms in thousands of cerebellar cells. Nature Biotechnology 36, 1197–1202 (2018).

23. Anvar, S. Y. et al. Full-length mRNA sequencing uncovers a widespread coupling between transcription initiation and mRNA processing. Genome Biology 19, 1–18 (2018).

24. Jing, Y. et al. Hybrid sequencing-based personal full-length transcriptomic analysis implicates proteostatic stress in metastatic ovarian cancer. Oncogene (2019).

25. Chen, H. et al. Long‐Read RNA Sequencing Identifies Alternative Splice Variants in Hepatocellular Carcinoma and Tumor‐Specific Isoforms. Hepatology hep.30500 (2019).

26. Tardaguila, M. et al. SQANTI: Extensive characterization of long-read transcript sequences for quality control in full-length transcriptome identification and quantification. Genome research (2018).

27. Ono, Y., Asai, K. & Hamada, M. PBSIM: PacBio reads simulatortoward accurate genome assembly. Bioinformatics 29, 119–121 (2012).

28. Stöcker, B. K., Köster, J. & Rahmann, S. SimLoRD: Simulation of long read data. Bioinformatics 32, 2704–2706 (2016).

29. Shcherbina, A. FASTQSim: Platform-independent data characterization and in silico read generation for ngs datasets. BMC research notes 7, 533 (2014).

30. Foulkes, W. D., Smith, I. E. & Reis-Filho, J. S. Triple-Negative Breast Cancer. New England Journal of Medicine 363, 1938–1948 (2010).

31. Kawazu, M. et al. Integrative analysis of genomic alterations in triple-negative breast cancer in association with homologous recombination deficiency. PLoS Genetics 13, 1–23 (2017).

32. Polak, P. et al. A mutational signature reveals alterations underlying deficient homologous recombination repair in breast cancer. Nature Genetics 49, 1476–1486 (2017).

33. Koboldt, D. C. et al. Comprehensive molecular portraits of human breast tumours. Nature 490, 61–70 (2012).

34. Nik-Zainal, S. et al. Landscape of somatic mutations in 560 breast cancer whole-genome sequences. Nature 534, 47–54 (2016).

35. Davies, H. et al. HRDetect is a predictor of BRCA1 and BRCA2 deficiency based on mutational signatures. Nature Medicine 23, 517–525 (2017).

36. McCann, K. E. & Hurvitz, S. A. Advances in the use of PARP inhibitor therapy for breast cancer. Drugs in Context 7, 1–30 (2018).

37. Chen, J. et al. Pervasive functional translation of noncanonical human open reading frames. Science (New York, N.Y.) 367, 1140–1146 (2020).

38. Garczyk, S. et al. AGR3 in breast cancer: Prognostic impact and suitable serum-based biomarker for early cancer detection. PloS one 10, e0122106 (2015).

39. Wali, V. B. et al. Identification and validation of a novel biologics target in triple negative breast cancer. Scientific reports 9, 14934 (2019).

40. Pampalakis, G. et al. The klk5 protease suppresses breast cancer by repressing the mevalonate pathway. Oncotarget 5, 2390–403 (2014).

41. Choi, S. K., Kim, H. S., Jin, T. & Moon, W. K. LOXL4 knockdown enhances tumor growth and lung metastasis through collagen-dependent extracellular matrix changes in triple-negative breast cancer. Oncotarget 8, 11977–11989 (2017).

42. Furuyama, K. et al. Continuous cell supply from a sox9-expressing progenitor zone in adult liver, exocrine pancreas and intestine. Nature genetics 43, 34–41 (2011).

43. Bjørklund, S. S. et al. Widespread alternative exon usage in clinically distinct subtypes of invasive ductal carcinoma. Scientific reports 7, 5568 (2017).

44. Gorello, P. et al. T(3;11)(Q12;P15)/NUP98-LOC348801 fusion transcript in acute myeloid leukemia. Haematologica 93, 1398–1401 (2008).

45. Qian, X. et al. The tensin-3 protein, including its sh2 domain, is phosphorylated by src and contributes to tumorigenesis and metastasis. Cancer cell 16, 246–58 (2009).

46. Cao, X. et al. A phosphorylation switch controls the spatiotemporal activation of rho GTPases in directional cell migration. Nature Communications 6, (2015).

47. Katz, M. et al. A reciprocal tensin-3-cten switch mediates egf-driven mammary cell migration. Nature cell biology 9, 961–9 (2007).

48. Sedlazeck, F. J. et al. Accurate detection of complex structural variations using single-molecule sequencing. Nature methods 15, 461–468 (2018).

49. Stephens, Z., Wang, C., Iyer, R. K. & Kocher, J.-P. Detection and visualization of complex structural variants from long reads. BMC bioinformatics 19, 508 (2018).

50. Carithers, L. J. et al. A novel approach to high-quality postmortem tissue procurement: The gtex project. Biopreservation and biobanking 13, 311–9 (2015).

51. Li, Y. et al. Patterns of somatic structural variation in human cancer genomes. Nature 578, 112–121 (2020).

52. Kim, P. & Zhou, X. FusionGDB: Fusion gene annotation database. Nucleic acids research 47, D994–D1004 (2019).

53. Mangeney, M. et al. Placental syncytins: Genetic disjunction between the fusogenic and immunosuppressive activity of retroviral envelope proteins. Proceedings of the National Academy of Sciences 104, 20534–20539 (2007).

54. Shuai, S. et al. The u1 spliceosomal RNA is recurrently mutated in multiple cancers. Nature 574, 712–716 (2019).

55. Suzuki, H. et al. Recurrent noncoding u1 snRNA mutations drive cryptic splicing in SHH medulloblastoma. Nature 574, 707–711 (2019).

56. Shen, S., Wang, Y., Wang, C., Wu, Y. N. & Xing, Y. SURVIV for survival analysis of mRNA isoform variation. Nature Communications 7, 1–11 (2016).

57. Silvester, J. et al. Gene isoforms as expression-based biomarkers predictive of drug response in vitro. Nature Communications 8, (2017).

58. Kahles, A. et al. Comprehensive analysis of alternative splicing across tumors from 8,705 patients. Cancer cell 34, 211–224.e6 (2018).

59. Manning, K. S. & Cooper, T. A. The roles of RNA processing in translating genotype to phenotype. Nature Reviews Molecular Cell Biology 18, 102–114 (2016).

60. Franco, H. L. et al. Enhancer transcription reveals subtype-specific gene expression programs controlling breast cancer pathogenesis. Genome research 28, 159–170 (2018).

61. Van den Berge, K., Soneson, C., Robinson, M. D. & Clement, L. StageR: A general stage-wise method for controlling the gene-level false discovery rate in differential expression and differential transcript usage. Genome biology 18, 151 (2017).

62. Shiraishi, Y. et al. An empirical bayesian framework for somatic mutation detection from cancer genome sequencing data. Nucleic acids research 41, e89 (2013).

63. Dobin, A. et al. STAR: Ultrafast universal rna-seq aligner. Bioinformatics (Oxford, England) 29, 15–21 (2013).

64. Patro, R., Duggal, G., Love, M. I., Irizarry, R. A. & Kingsford, C. Salmon provides fast and bias-aware quantification of transcript expression. Nature methods 14, 417–419 (2017).

65. Li, B. & Dewey, C. N. RSEM: Accurate transcript quantification from rna-seq data with or without a reference genome. BMC bioinformatics 12, 323 (2011).

66. Li, Y. I. et al. Annotation-free quantification of RNA splicing using LeafCutter. Nature Genetics 50, 151–158 (2018).

67. Tardaguila, M. et al. SQANTI: extensive characterization of long-read transcript sequences for quality control in full-length transcriptome identification and quantification. Genome Research 28, 396–411 (2018).

68. Camacho, C. et al. BLAST+: Architecture and applications. BMC bioinformatics 10, 421 (2009).

69. Byrne, A., Cole, C., Volden, R. & Vollmers, C. Realizing the potential of full-length transcriptome sequencing. Philosophical transactions of the Royal Society of London. Series B, Biological sciences 374, 20190097 (2019).

70. Anvar, S. Y. et al. Full-length mRNA sequencing uncovers a widespread coupling between transcription initiation and mRNA processing. Genome Biology 19, (2018).

71. Frazee, A. C., Jaffe, A. E., Langmead, B. & Leek, J. T. Polyester: Simulating rna-seq datasets with differential transcript expression. Bioinformatics (Oxford, England) 31, 2778–84 (2015).

72. Love, M. I., Huber, W. & Anders, S. Moderated estimation of fold change and dispersion for rna-seq data with deseq2. Genome Biology 15, 550 (2014).

73. Love, M. I., Soneson, C. & Patro, R. Swimming downstream: Statistical analysis of differential transcript usage following salmon quantification. F1000Research 7, 952 (2018).

74. Soneson, C., Love, M. I. & Robinson, M. D. Differential analyses for rna-seq: Transcript-level estimates improve gene-level inferences. F1000Research 4, (2015).

75. Anders, S., Reyes, A. & Huber, W. Detecting differential usage of exons from rna-seq data. Genome Research 22, 4025 (2012).

76. Nowicka, M. & Robinson, M. D. DRIMSeq: A dirichlet-multinomial framework for multivariate count outcomes in genomics [version 2; referees: 2 approved]. F1000Research 5, (2016).

77. Trincado, J. L. et al. SUPPA2: Fast, accurate, and uncertainty-aware differential splicing analysis across multiple conditions. Genome biology 19, 40 (2018).

78. Saraiva-Agostinho, N. & Barbosa-Morais, N. L. Psichomics: Graphical application for alternative splicing quantification and analysis. Nucleic Acids Research 47, e7 (2019).

79. Finn, R. D., Clements, J. & Eddy, S. R. HMMER web server: Interactive sequence similarity searching. Nucleic Acids Research 39, W29–W37 (2011).

80. El-Gebali, S. et al. The pfam protein families database in 2019. Nucleic Acids Research 47, D427–D432 (2018).

81. Oki, S. et al. ChIP-atlas: A data-mining suite powered by full integration of public chip-seq data. EMBO reports 19, (2018).

