## SupplementaryFigures for "Multi-sample Full-length Transcriptome Analysis of 22 Breast Cancer Clinical Specimens with Long-Read Sequencing"

### Supplementary Notes

###### Simulations in various conditions

As for expression data, MuSTA acquires transcript per million (TPM) from shot-read RNAseq, and PBcount from SMRT sequencing. We investigated which two, if not both are adequate to be used for DTU detection. The detection performance was the greatest when using both in combination to detect DTU, with true positive rate (TPR) of 0.95 and false discovery rate (FDR) of 0.059 when the target FDR was set to 0.01 (Supplementary Fig. 2a). In transcript level, detection performance was not satisfactory compared to gene level (TPR was 0.89 and FDR was 0.32).

Because IsoSeq is able to detect a large number of unannotated isoforms, we also calculated TPR and FDR for genes with more than 15 isoforms (Supplementary Fig. 2b). Although there was no significant change in sensitivity, specificity for those genes was significantly lower than that for all genes. We note that we excluded accidentally detected unannotated genes when calculating TPR and FDR for genes with more than 15 isoforms, and it was likely to be responsible for TPR increase.

Previously, it has been reported that in the detection of DTU, exclusion of isoforms with lower expression improves performance^1^. Therefore, we’ve tried two kinds of pre-filtering. First, we’ve found the sensitivity to be lower when we filtered out isoforms with low short-read coverage (post-alignment prefiltering) (Supplementary Fig. 2c). Second, a larger number of isoforms per gene could result in increased miss-aligned short-read fragments; therefore, we experimented with only isoforms with substantial PBcount (major isoforms) for obtaining short-read TPM (pre-alignment prefiltering). Although DTU detected in major isoforms had better specificity than those detected in full isoforms, the sensitivity was lower. In conclusion, we yielded the best performance when not using pre-filtering, according to our simulation. We hypothesized that this result was due to the method of simulations, where we assigned DTU independently from isoform expression. Biologically valuable DTU should have relatively high expression among isoforms that belong to the same genes. Therefore, we examined the effect of prefiltering in the simulations based on the permutation of the breast cancer dataset, where we assigned DTU to isoforms with the two highest expression (Methods). Nevertheless, we observed that prefiltering substantially lowered TPR regardless of NIC rates against all DTU isoforms (Supplementary Fig.4). Our results implicate that, although we didn’t examine the most suitable parameters for prefiltering, DTU inference was best performed without prefiltering when using a MuSTA-derived transcriptome.

Next, we looked into the performance of DTU detection according to four conditions: (i) the number of samples of each group, (ii) fold change of DTU isoforms, (iii) the total short-read number, and (iv) the total long-read number. Sensitivity was consistently high regardless of the sample number of each group (Supplementary Fig. 2d). As the fold change in DTU increase, FDR for DTU detection in transcript level increased unexpectedly (Supplementary Fig. 2e). As the total short-read number increase, FDR for DTU detection in transcript level again increased unexpectedly (Supplementary Fig. 2f). These contradictory declines in specificity were associated with the increase in the total short-read count and the difference between groups in the short-read count of isoforms not defined as DTU (Supplementary Fig. 3a-d). On the other hand, TPR showed little change when long-read sequencing depth increased (Supplementary Fig. 2f). This may reflected that the increase in the number of isoforms detected in long-read RNAseq was limited (Supplementary Fig. 3e). This number seemed to be saturated at around 180,000, reflecting the setting that all simulated transcripts were generated from 203,673 known transcripts in GENCODE. This situation is unrealistic, and indeed the number of isoforms was not saturated when MuSTA was applied to breast cancer specimens (Figure 3b).

###### Relationship between short-read expression and long-read detection and expression

We studied the relationship between short-read and long-read expression using TN2 as a representative sample. TPM calculated by salmon and TPM calculated by STAR and RSEM matched well in gene level, however, in transcript level, the correlation was weaker (Supplementary Fig. 4a). In some samples, calculation with STAR and RSEM was not completed in our computational environment because they required more than 128 gigabytes. TPM of all isoforms and TPM of (pre-filtered) major isoforms had a very strong correlation in gene level, however the correlation was weaker in transcript level (Supplementary Fig. 4b). Almost all major isoforms showed higher amount of TPM, possibly because the short-reads derived from excluded isoforms was inappropriately aligned on them. Short-read TPM and PBcount had a weak positive correlation. Comparing the correlation in gene level and transcript level, the latter had an even weaker correlation (Supplementary Fig. 4c-e).

###### Alternative Splicing in MuSTA-transcriptome

We examined alternative splicing in the transcriptome. There, we found that mutually exclusive exons were of the largest number, and single exon skipping/insertion showed the smallest number (Supplementary Fig. 5a, b). Possibly due to the high sensitivity of IsoSeq for detection of isoforms, there is a high probability to detect multiple exon skipping events for genes with a large number of isoforms. Using psichomics^2^ and conducting principal component analysis about alternative splicing events, first principal component (PC) reflected subtypes well (Supplementary Fig. 5c). Second PC expressed heterogeneity in TNBC, and specifically TNM1 showed different value compared to others. TNM1 is TNBC with BRCA1 methylation, and it is unknown why it became to be an outlier. Molecularly association / molecularly and mutually exclusivity between distal exons (dMAP/dMEP) are possible to detect with long-read sequencing, and dMAP had 1250 events (831 genes) while dMEP had 847 events (596 genes).

###### Comparison of DGE and DTU between GENCODE-transcriptome and MuSTA-transcriptome

The methods to detect the genes with DGE or DTU with GENCODE-transcriptome were compared with those with MuSTA-transcriptome (Supplementary Fig. 10a-c). The number of isoforms per GENCODE gene fell between 1 and 378 (median 9) in MuSTA-transcriptome. Only 0.97% of the genes had more than 100 isoforms. While more than half of the identified DGE genes (n=3014) were detected with both methods, the DTU genes identified with both methods (n=222) were 17% and 48% of the DTU genes identified with GENCODE (n=1307) and MuSTA-transcriptome (n=465), respectively, although those DTU genes were mostly present in both annotations. DTU isoforms detected in both methods (n=80) accounted for only 5.2% of all the DTU isoforms in GENCODE (n=1546), and 16% in MuSTA-transcriptome (n=500). These higher rates in MuSTA-transcriptome support the conservative behavior of DTU inference with MuSTA-transcriptome.

### Supplementary Figures

###### Supplementary Figure 1

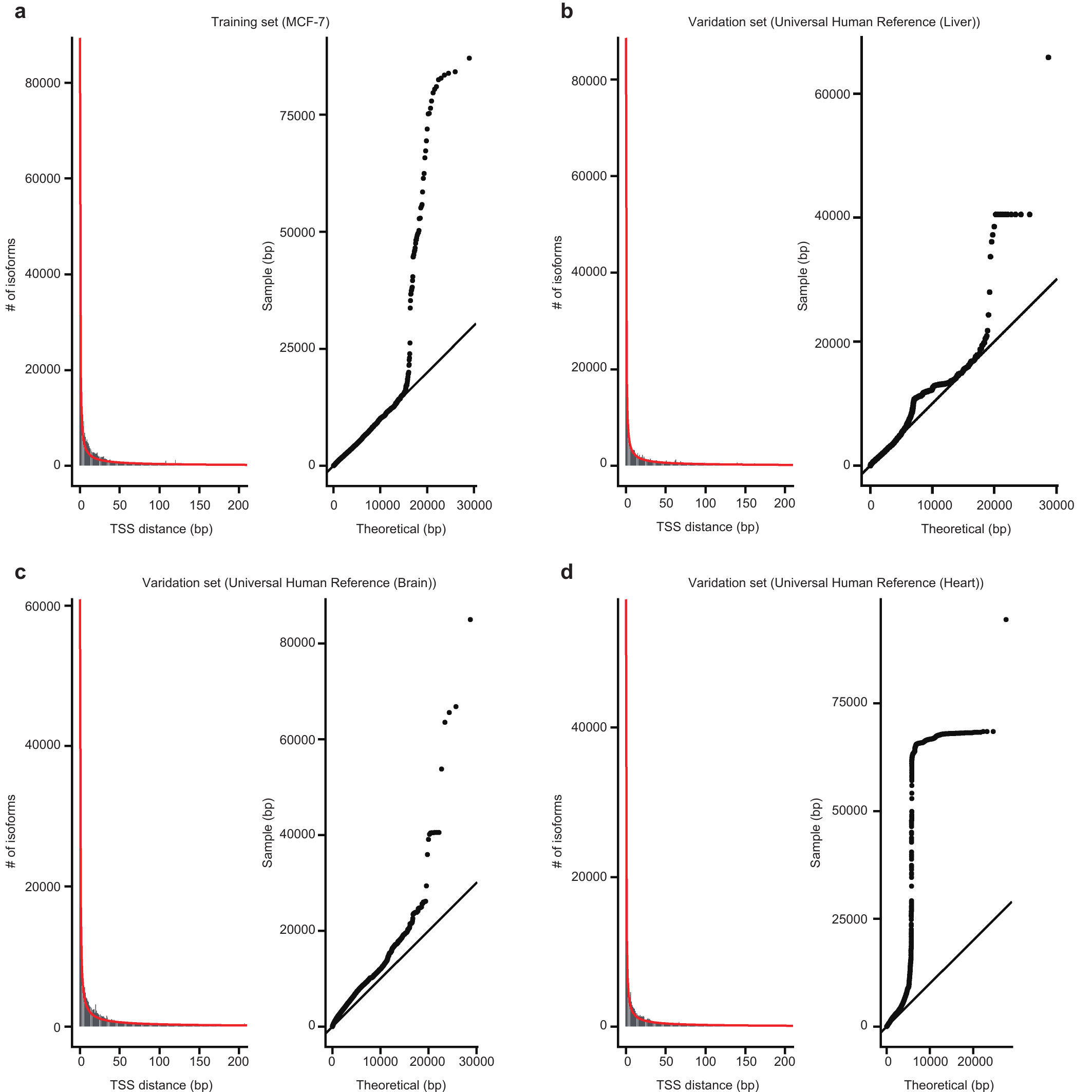

**Parameter estimation for RNA 5’ degradation.** Figure **a** shows training data set, and figures **b-d** show validation data sets. Left: The distance between TSS of FLNC read and nearest upstream TSS in GENCODE TSS. The red line shows fitted curve. Right: quantile-quantile plot of curve fitting. TSS, transcript start site; FLNC, full-length non-chimeric.

###### Supplementary Figure 2

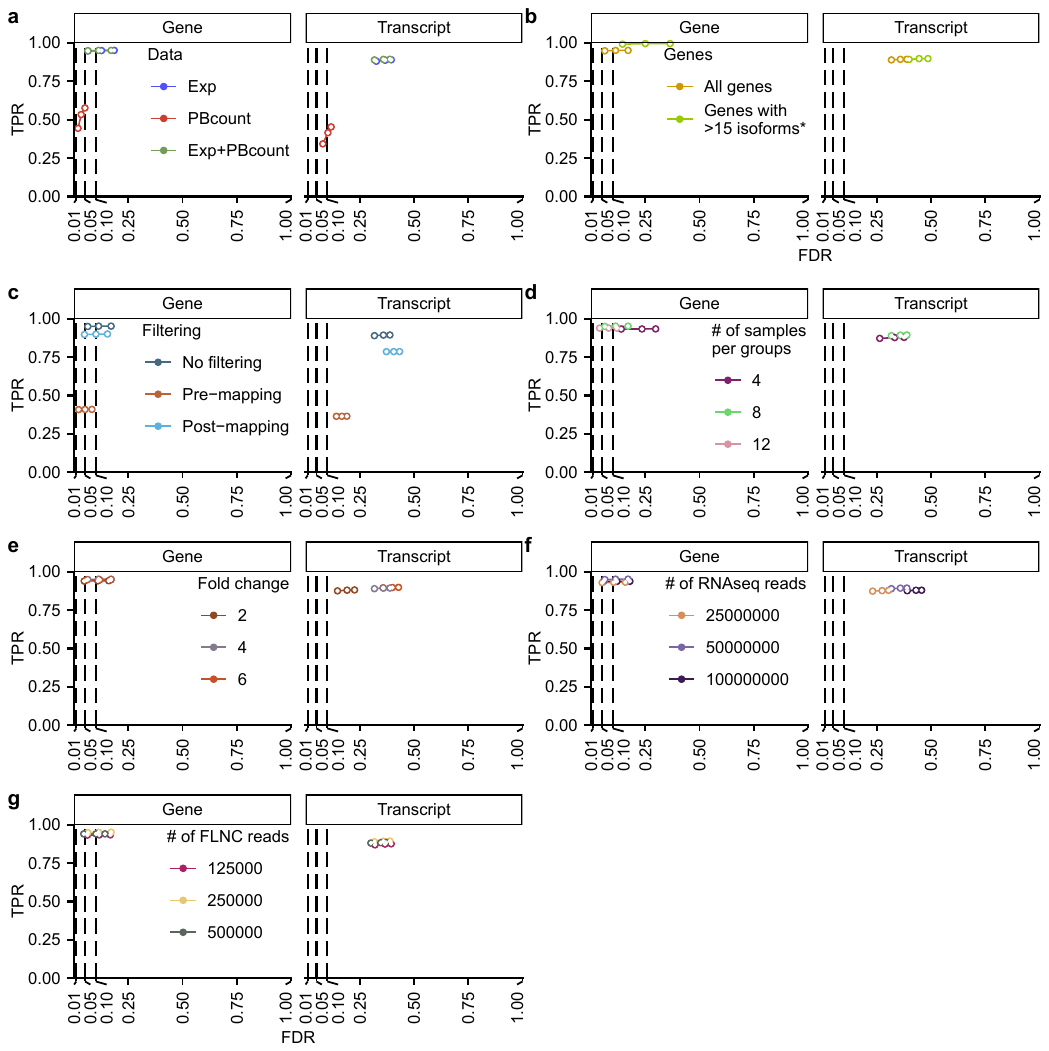

**True positive rate (TPR)-false discovery rate (FDR) plots of differential transcript usage (DTU) simulations.** The dots on each plot indicate the target FDR of 0.01, 0.05, and 0.1. Conditions were changed according to expression data (**a**), number of isoforms (**b**), isoform prefilter (**c**), number of samples in each group (**d**), fold change of DTU isoforms (**e**), and depth of short-read (**f**) and long-read (**g**) RNA sequencing. Note that we excluded accidentally detected unannotated genes when calculating TPR and FDR for genes with more than 15 isoforms (asterisk).

###### Supplementary Figure 3

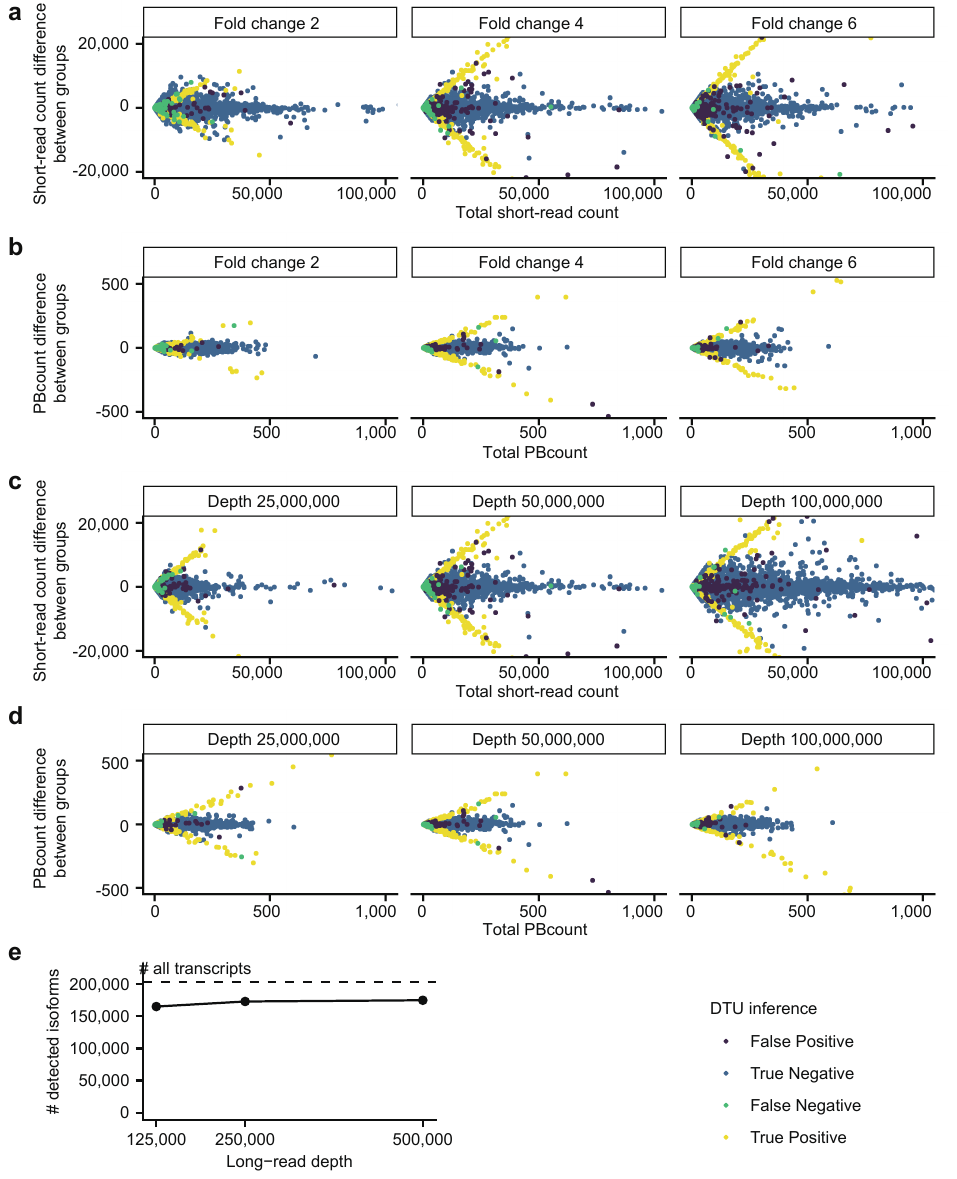

**The detailed simulation results under different condition.** **a-d**, The sum and the difference in expression between groups when (**a** and **b**) changing the fold change and (**c** and **d**) changing depth of short read. In **a** and **c**, expression data were short-read count, and in **b** and **d**, they were PBcount. **e**, The number of isoforms detected according to the depth of long-read.

###### Supplementary Figure 4

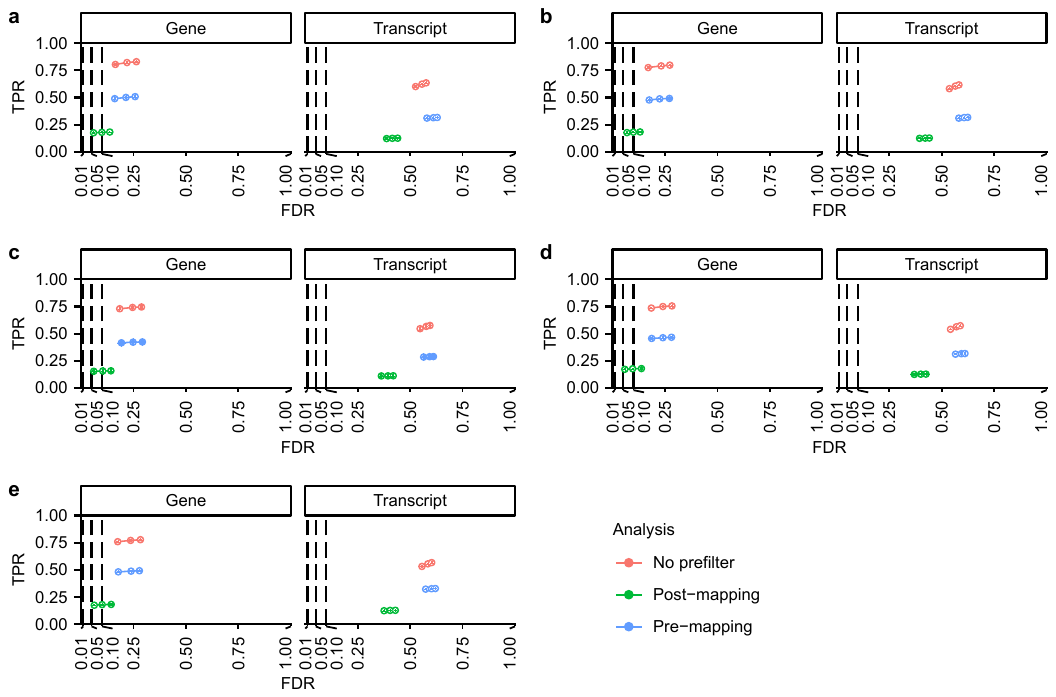

**TPR-FDR plots of DTU simulations based on the permutation of the breast cancer dataset.** NIC rates against all DTU isoforms were set to 0 (**a**), 0.25 (**b**), 0.5 (**c**), 0.75 (**d**), and 1 (**e**). The dots represent the mean and the error bars represent the standard error of three independent simulations.

###### Supplementary Figure 5

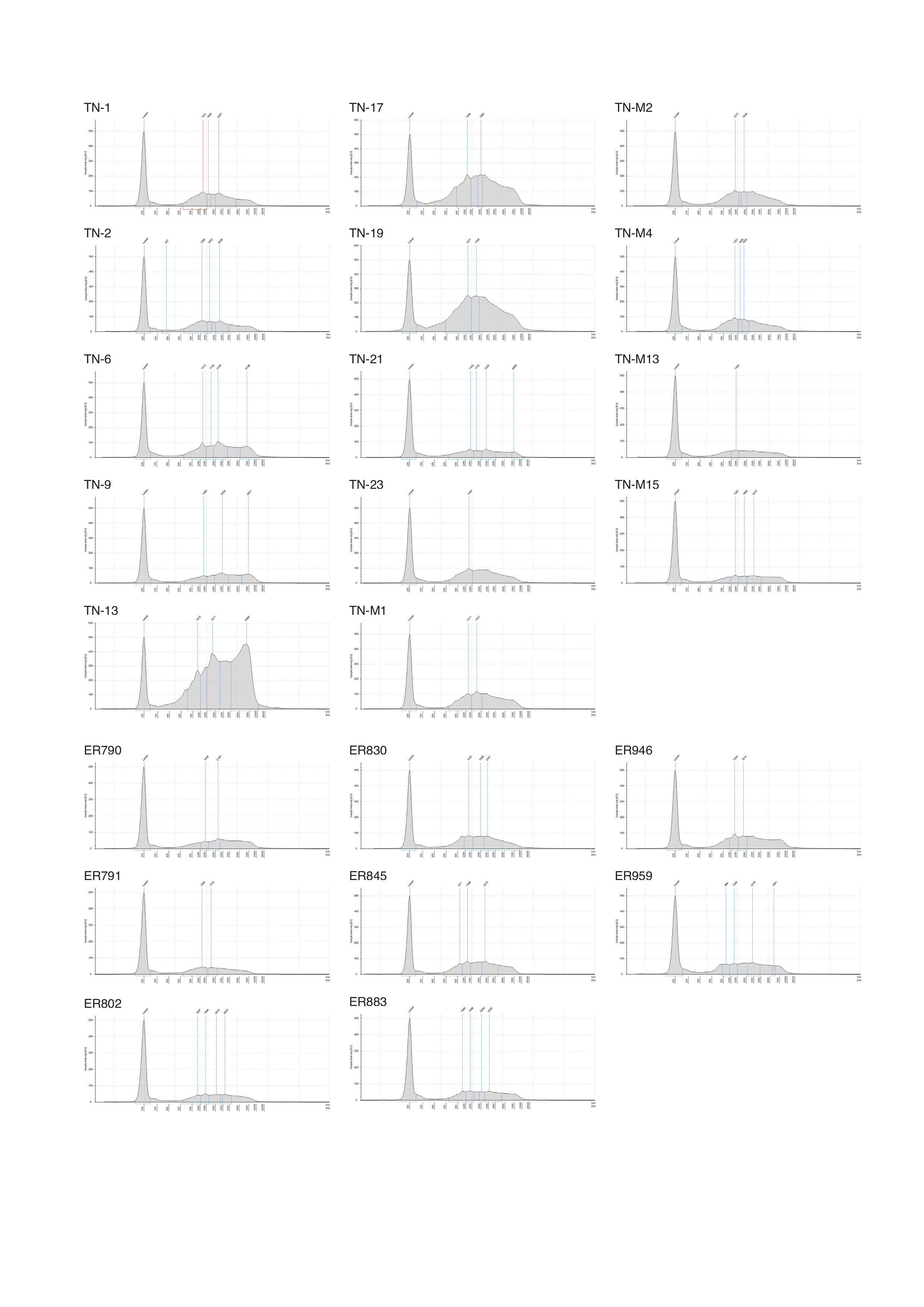

**The distribution of the read length resulted from SMRT sequencing of 22 breast cancer clinical specimens.**

###### Supplementary Figure 6

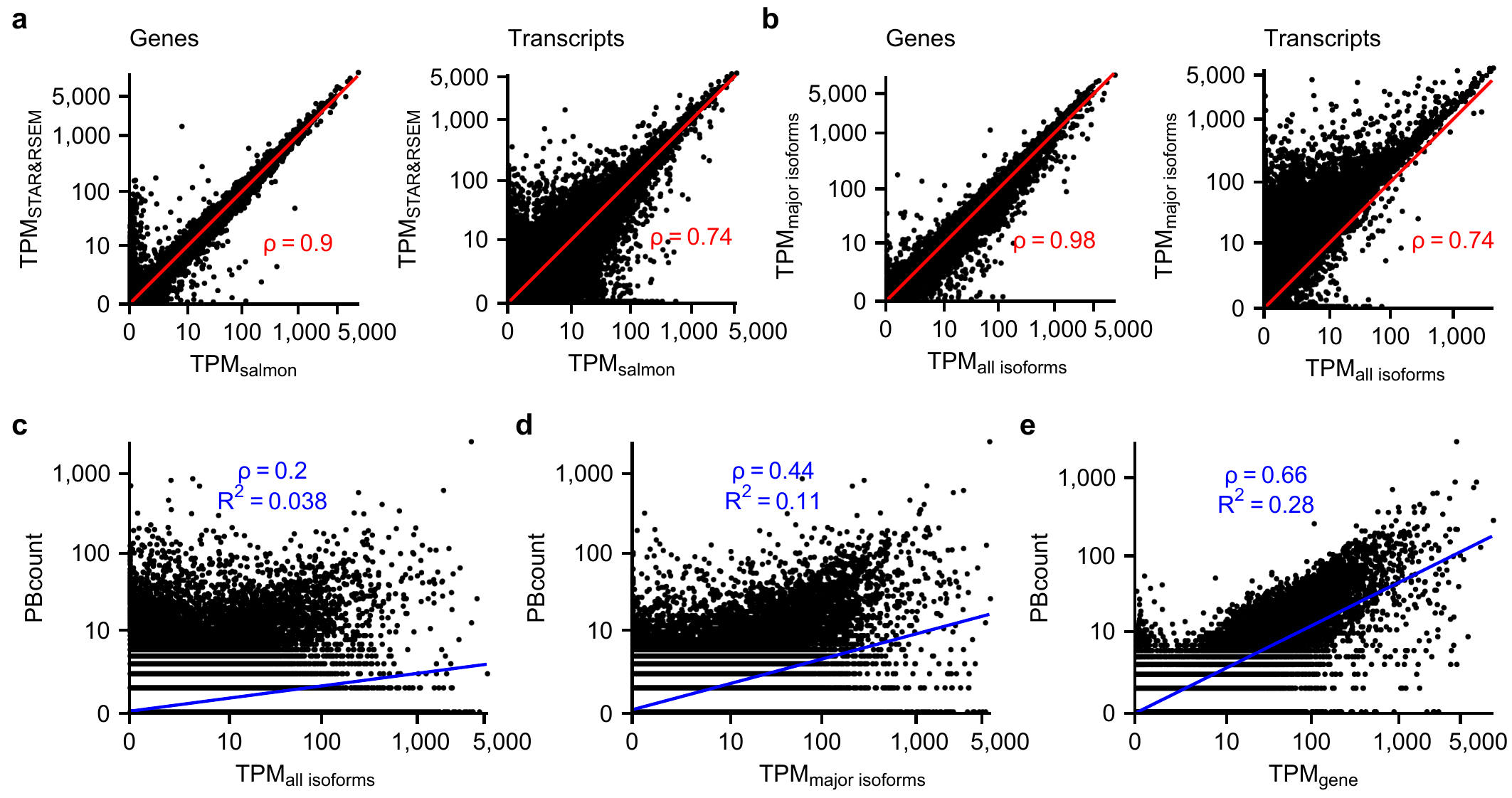

**Relationship between short-read TPM, long-read detection, and PBcount.** **a** and **b**, Difference of TPM in TN2, one of the TNBC samples, between different alignment methods (**a**) and with/without the pre-alignment isoform prefilter (**b**). Each dot represents a gene (left) or a transcript (right). Red lines indicate $y=x$, and $\rho$ means Spearman’s $\rho$. **c-e**, Relationship between PBcount and TPM in TN2 sample. **c** and **d** are in transcript-level, and **e** is in gene-level. $\rho$, Spearman’s $\rho$; $R^{2}$, R-squared value.

###### Supplementary Figure 7

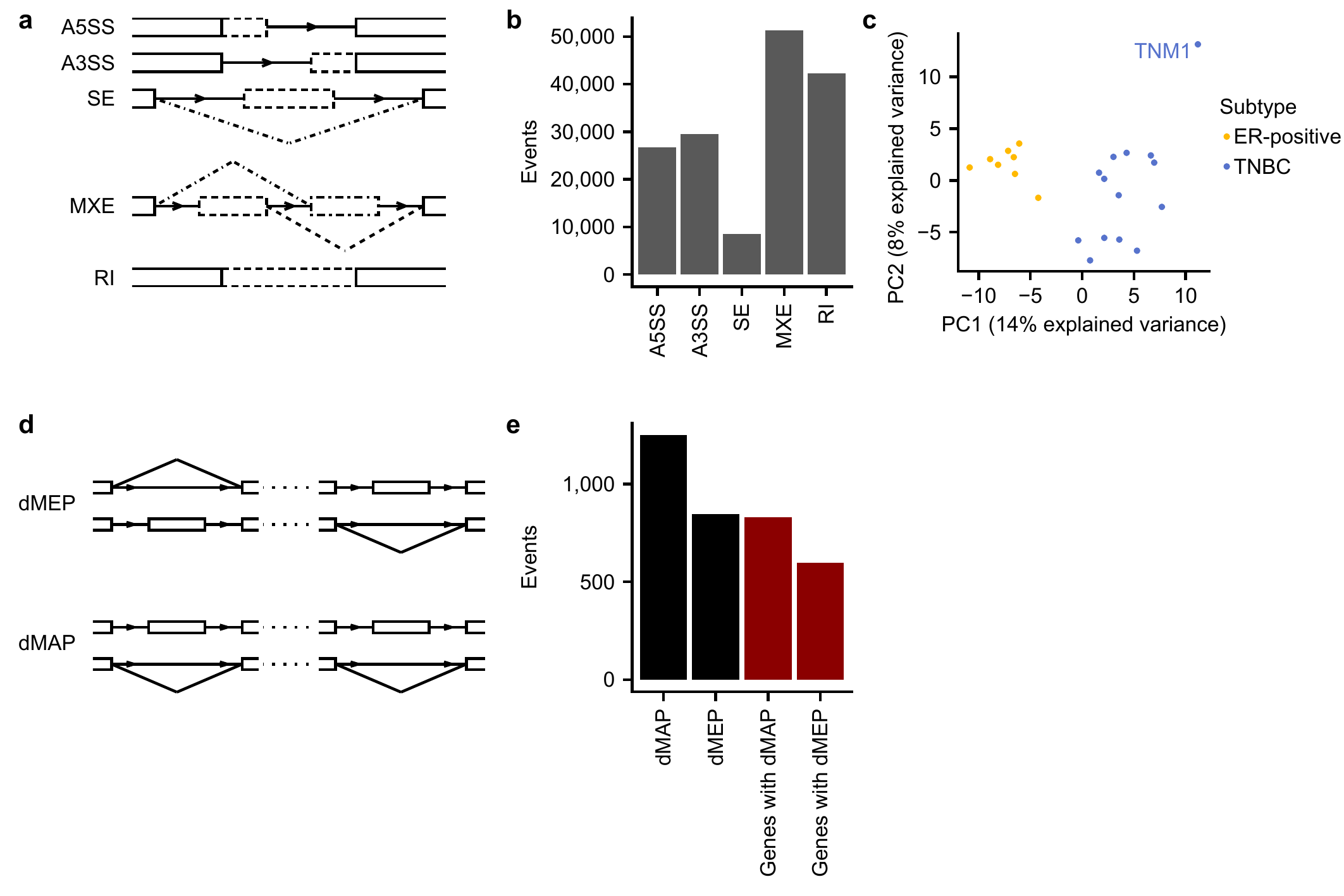

**Alternative splicing in 22 breast cancer clinical specimens.** **a**, A schema of alternative splicing categories. A5SS, alternative 5’ splice site; A3SS, alternative 3’ splice site; SE, single exon skipping/inclusion; MXE, mutually exclusive exons; RI, retained intron. **b**, Alternative splicing events across MuSTA-transcriptome from 22 breast cancer specimens. **c**, A principal component analysis using alternative splicing events. Axes represent first and second principal components. **d**, A schema of distal coupling between alternative splicing events. dMAP, distal molecularly associated exon pairs; dMEP, distant molecularly and mutually exclusive pairs. **e**, Bar plots showing the number of distal alternative splicing coupling.

###### Supplementary Figure 8

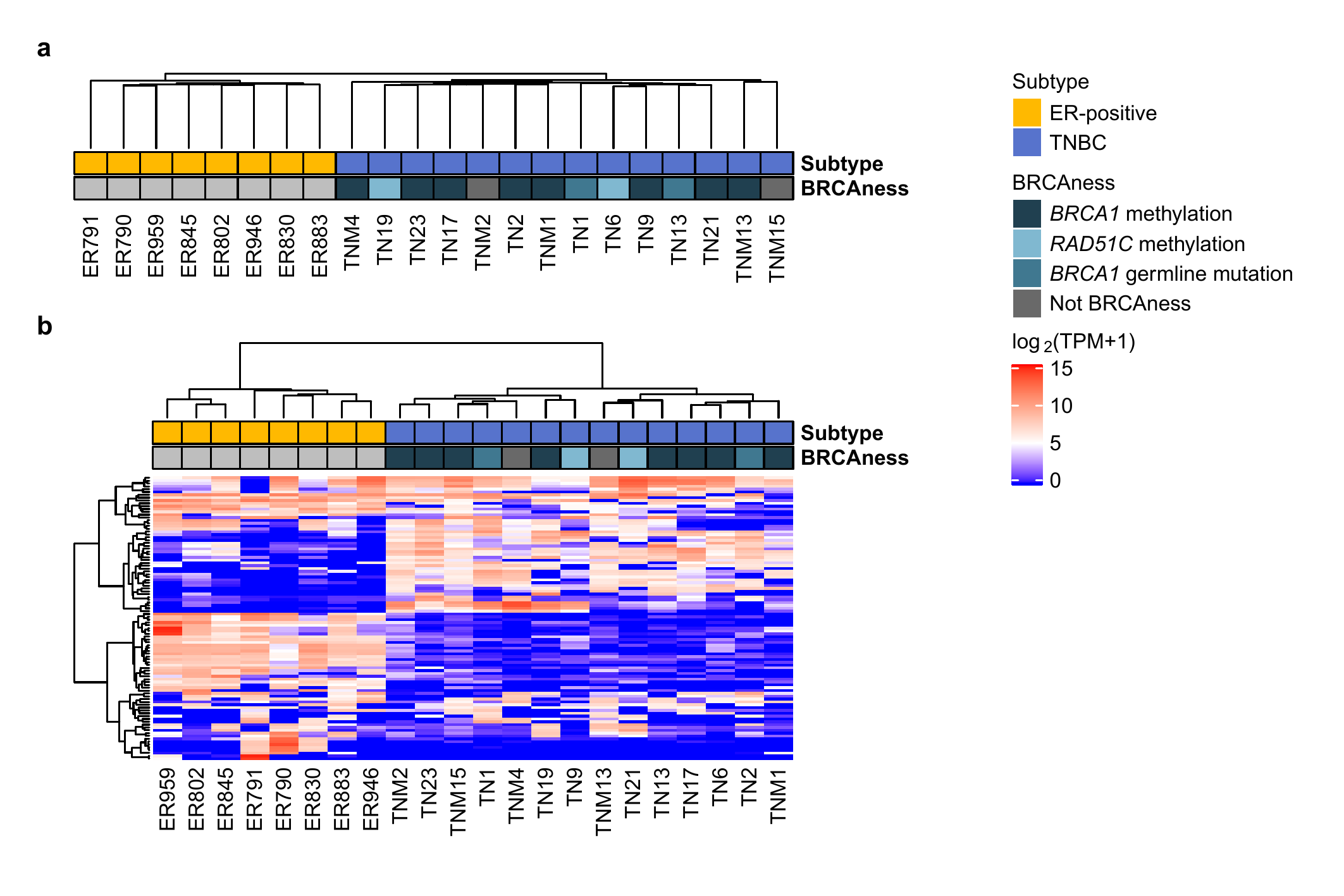

**Clustering reflecting subtypes, but not BRCAness.** Hierarchical clustering with isoform detection in MuSTA (**a**) and TPM of 100 isoforms with the highest standard deviation of TPM (**b**). In **a**, Samples turned out to separate into two groups which exactly matched with the breast cancer subtypes. BRCAness didn’t create a grouped cluster, and that was the same when TPM was used to make a regular clustering (**b**). Of note, detection results are not necessarily independent from isoform expression because isoforms with low expression are hard to detect using SMRT sequencing.

###### Supplementary Figure 9

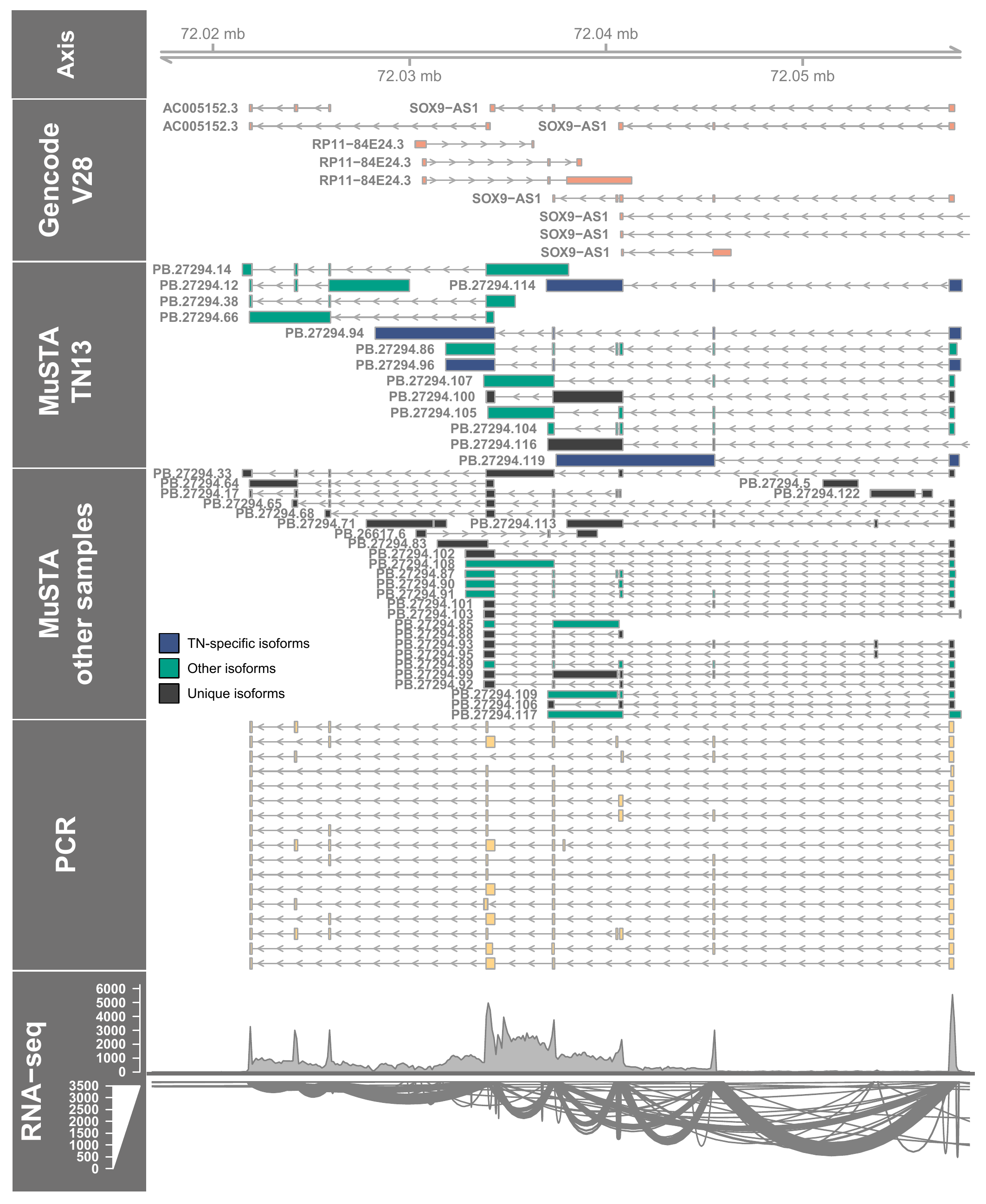

**Nested PCR validation of *SOX9-AS1/AC005152.3* readthrough transcripts.** From top to bottom, the areas represent genome axis, GENCODE annotation around *SOX9-AS1* and *AC005152.3*, MuSTA isoforms detected in TN13 sample, MuSTA isoforms detected in samples other than TN13, nested PCR products in TN13, and coverage and sashimi-plot of RNAseq in TN13, accordingly.

###### Supplementary Figure 10

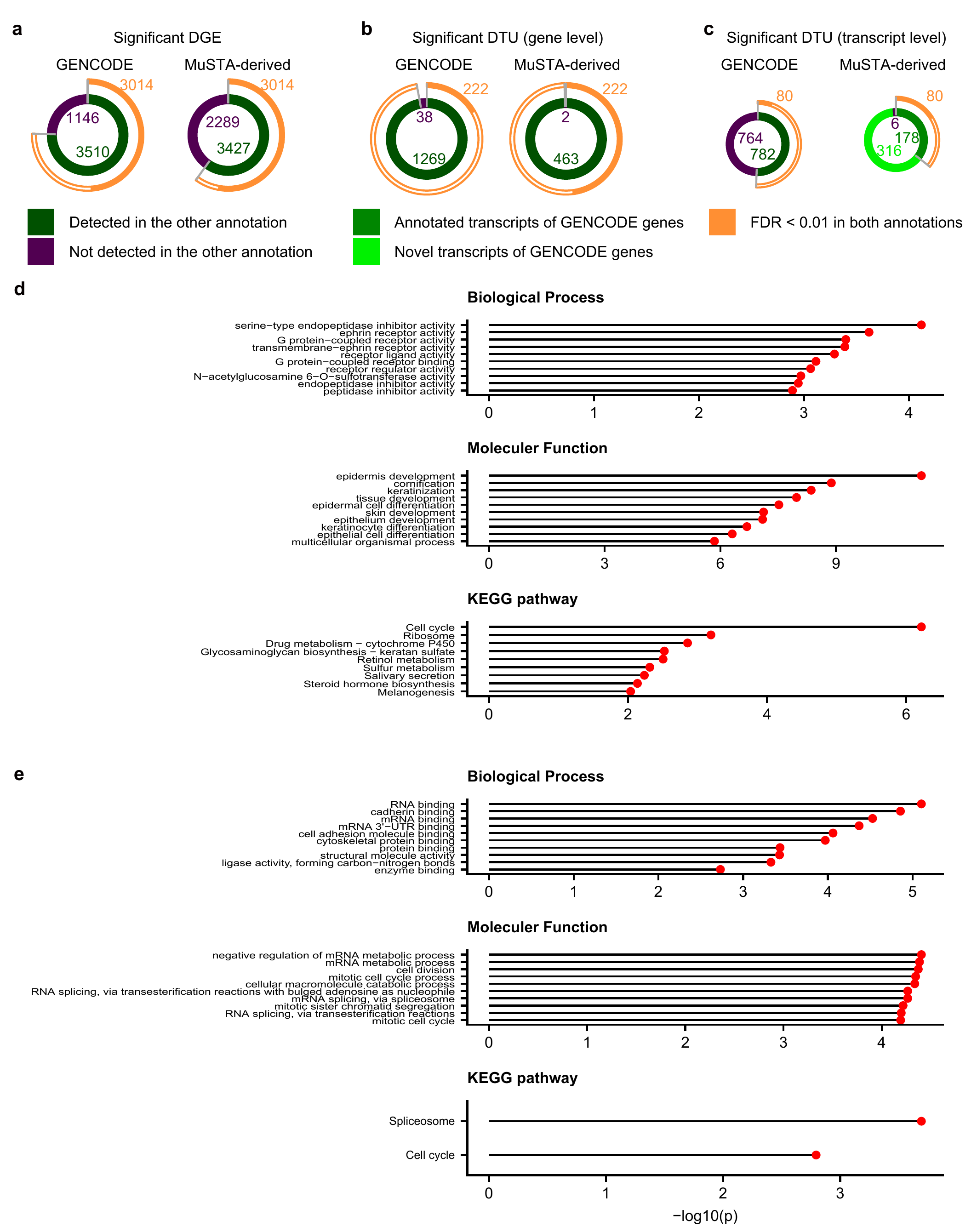

**Composition of DGE and DTU genes, and gene ontology and KEGG pathway enrichment analysis in DGE and DTU genes.** **a-c**, Donut plots representing genes with DGE (**a**), genes with DTU (**b**), and isoforms with DTU (**c**). Left, DGE genes, DTU genes, and DTU isoforms detected with the GENCODE transcriptome. Right, these detected with the MuSTA-derived transcriptome. Outer rings show the proportion of genes or isoforms which were labeled as DGE / DTU in both annotations with FDR < 0.01. The numbers represent the number of genes or isoforms in each group. **d** and **e**, Gene enrichment analysis for DGE (**d**) genes and DTU (**e**) genes for the MuSTA-derived transcriptome. P-values were calculated by hypergeometric test implemented in GOstats R package^3^. Top, biological process of gene ontology; center, molecular function of gene ontology; bottom, KEGG pathway.

###### Supplementary Figure 11

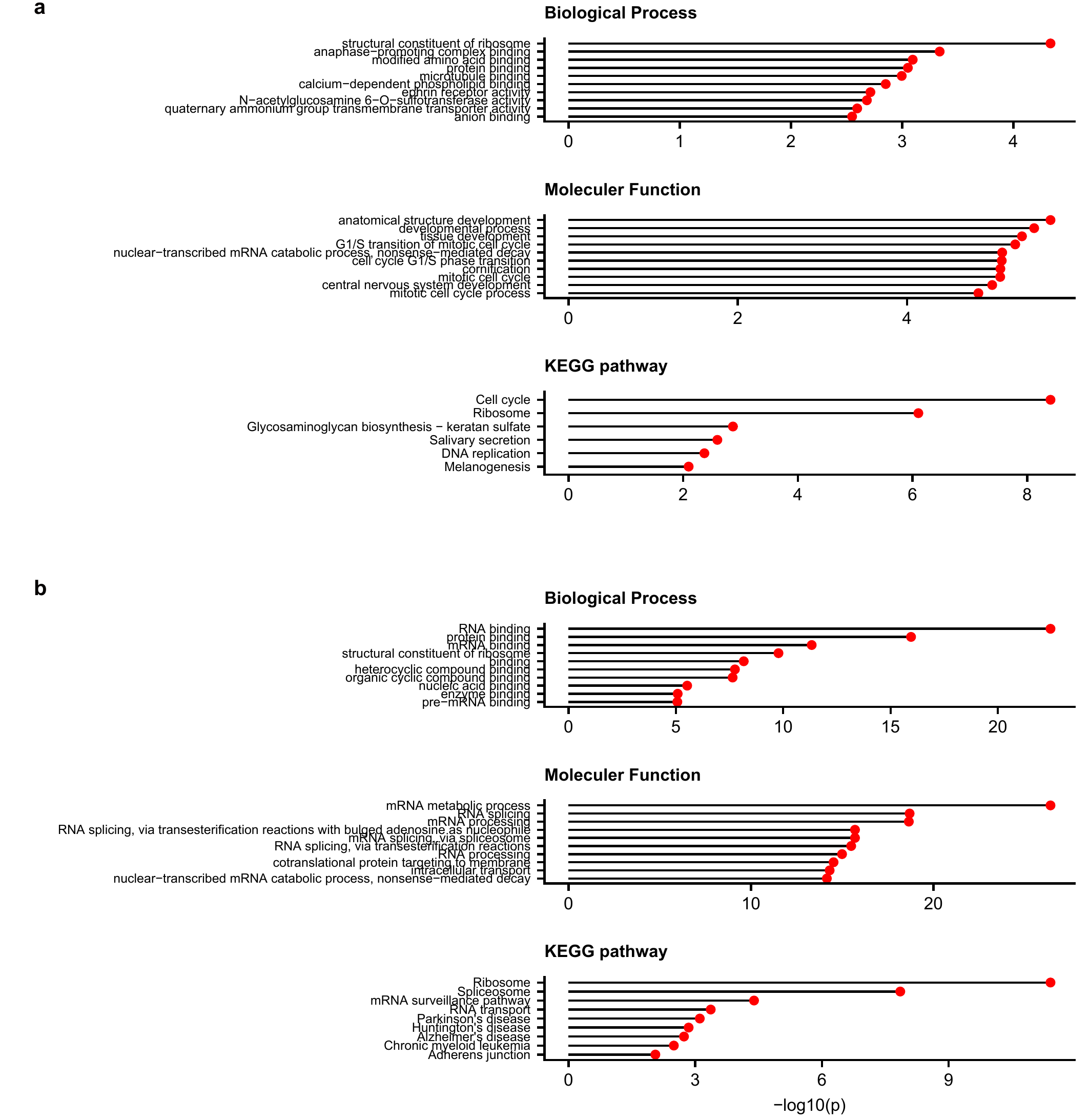

**Gene ontology and KEGG pathway enrichment analysis in DGE and DTU genes for the GENCODE annotation.** Gene enrichment analysis for DGE (**a**) genes and DTU (**b**) genes for the GENCODE transcriptome.

###### Supplementary Figure 12

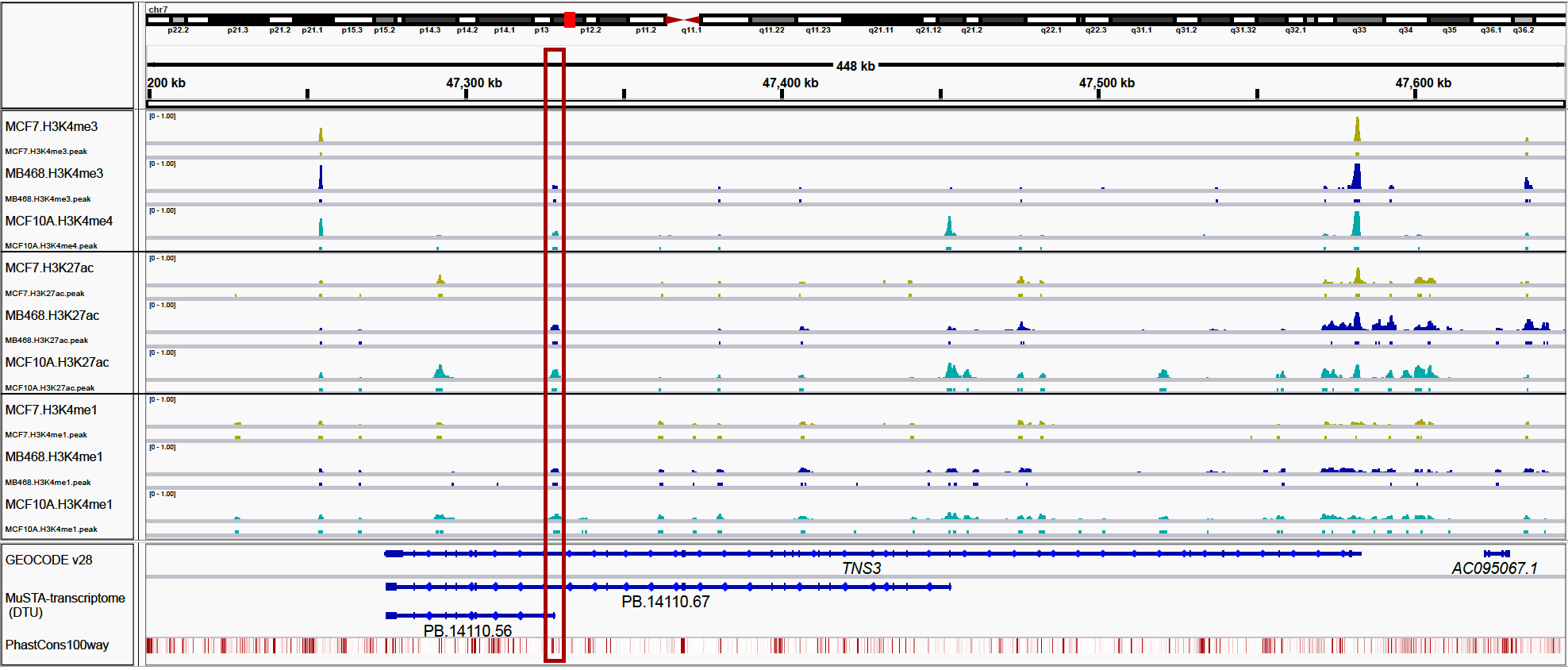
 **Snapshots of Integrative Genomics Viewer showing chromatin modifications of three breast cell lines and evolutionary conservation in *TNS3* region.** Shown were three chromatin modifications, H3K4me3 (enriched at promoters), H3K27ac1 and H3K4me1 (enriched at enhancers) in three breast normal/cancer cell lines, MCF-7 (ER-positive breast cancer), MDA-MB-468 (TNBC), and MCF-10A (normal breast epithelium).

###### Supplementary Figure 13

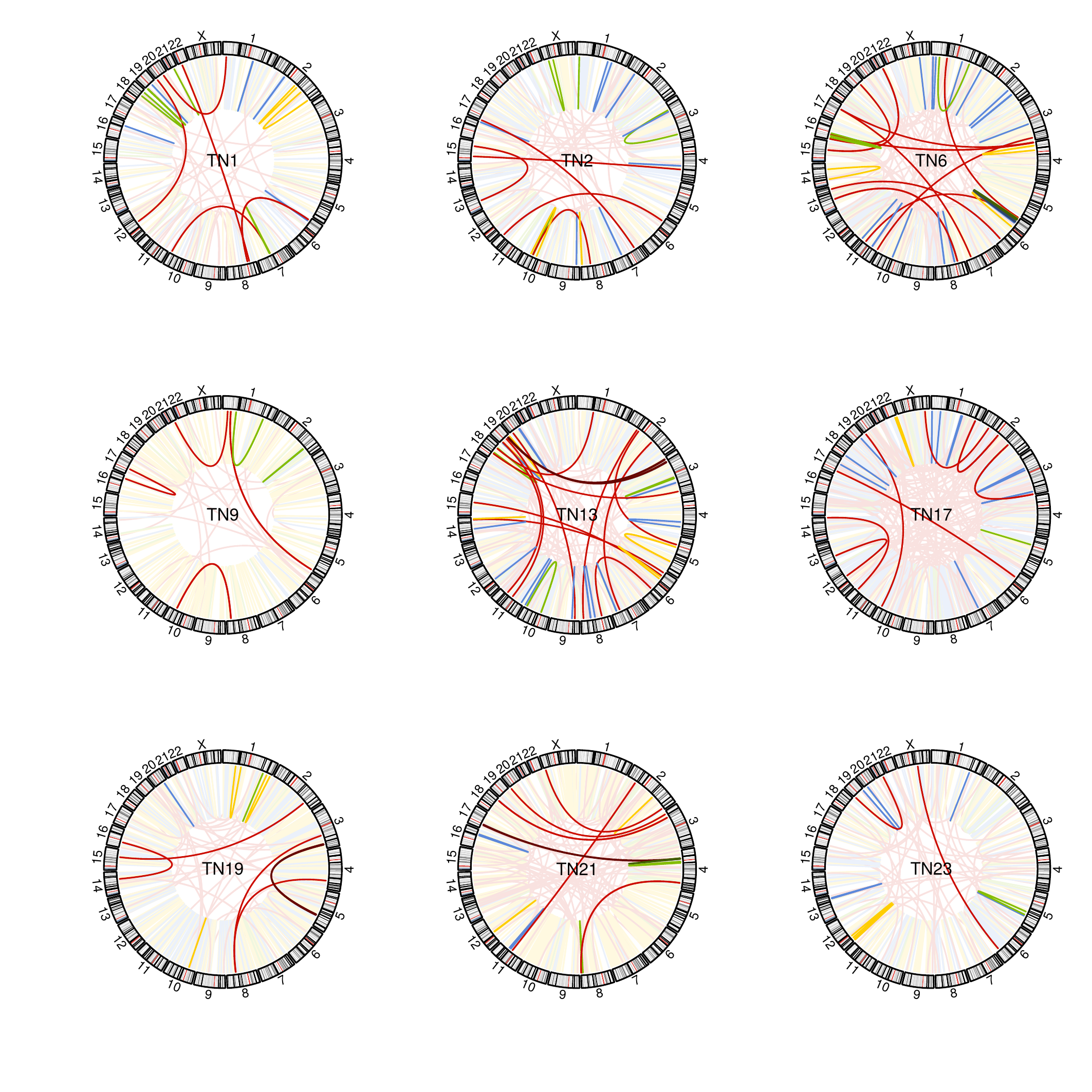

**Circos plots of structural variants and fusion transcripts.** Colored lines and shaded lines represent structural variants (SV) with or without corresponding fusion transcripts, respectively. Colors correspond to SV types, i.e. yellow to deletion, green to inversion, blue to tandem duplication, and red to translocation. Colors are darkened for nested SV with fusion transcripts.

###### Supplementary Figure 14

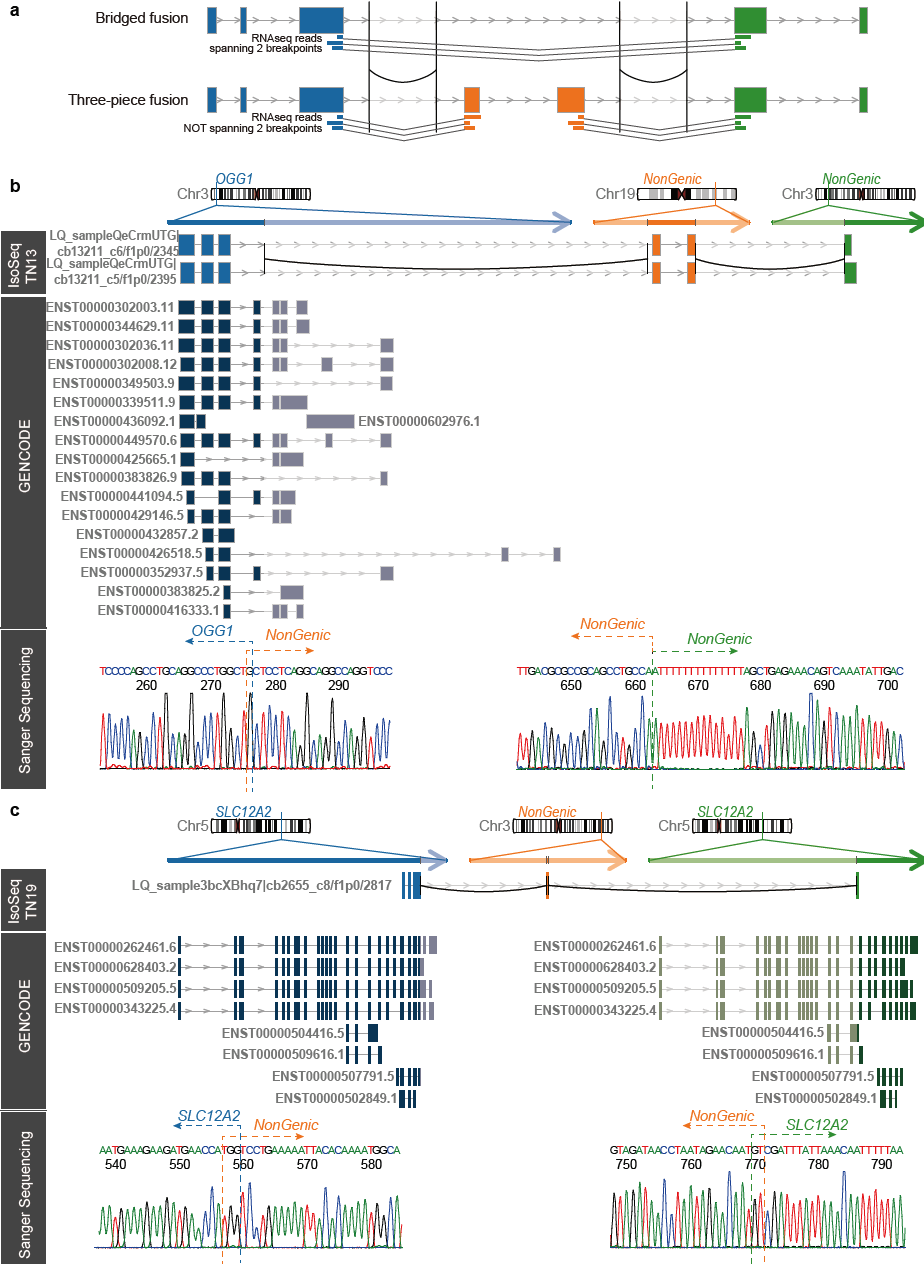

**PCR-validated three-piece fusion transcripts other than figure 5.** **a**, Schematic image of difference between bridged fusion transcript and three-piece fusion transcript. **b and c**, Structure of three-piece fusion transcripts from *OGG1-NonGenic-NonGenic* (**b**) and *SLC12A2-NonGenic-SLC12A2* (**c**). The genomic axes represent three original genomic regions. Below them are chimeric IsoSeq cluster reads. Curves correspond to structural variants detected with whole genome sequencing data. The category “GENCODE” shows annotated transcripts. Outside regions of structural variants are shaded. Exon-intron structures don’t necessarily reflect accurate length for visibility.

###### Supplementary Figure 15

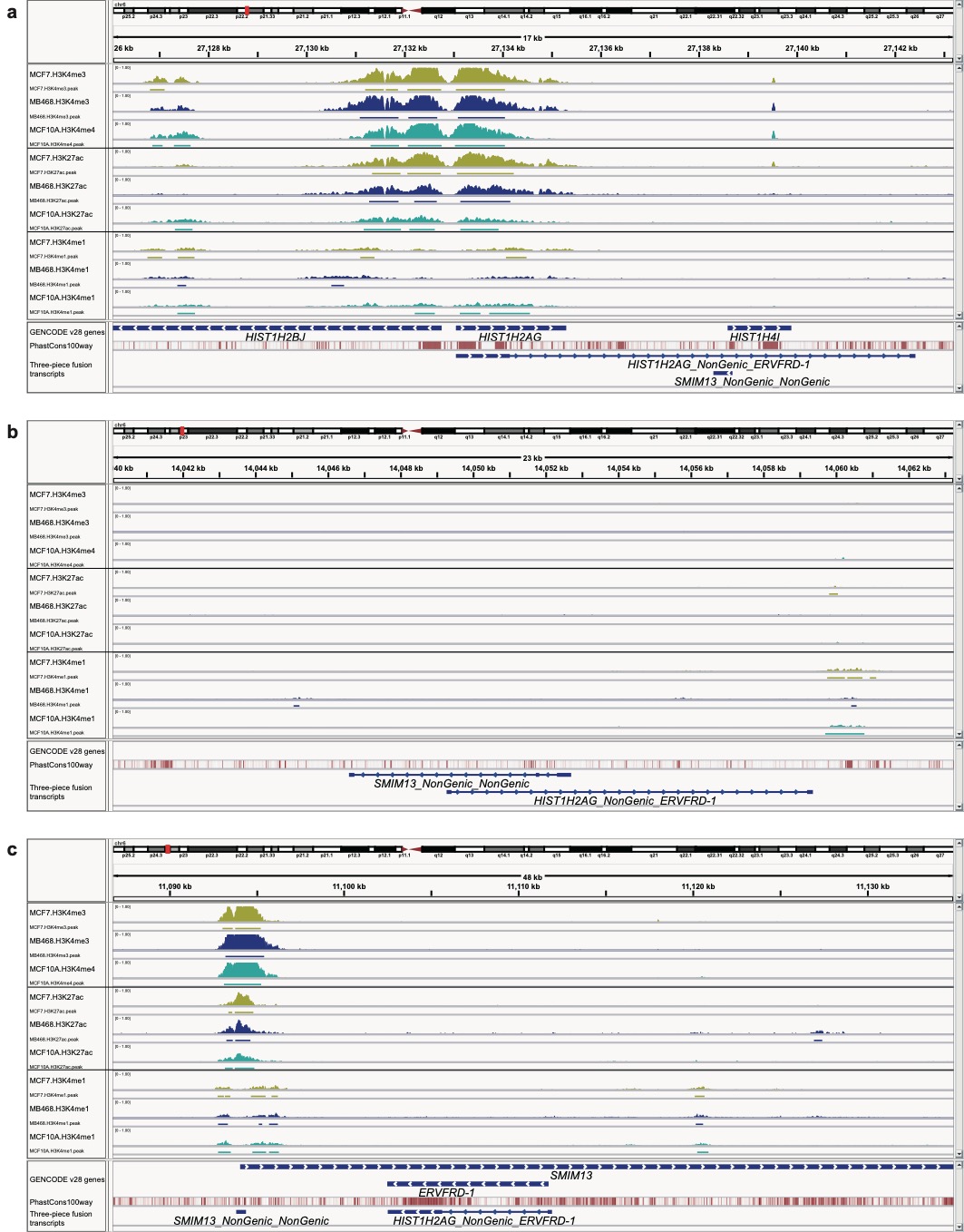

**Chromatin modifications related to *HIST1H2AG-NonGenic-ERVFRD-1.*** Snapshots of Integrative Genomics Viewer in the three regions where *HIST1H2AG* (**a**), *NonGenic* (**b**), and *ERVFRD-1* (**c**) fragments of the bridged fusion transcript, *HIST1H2AG-NonGenic-ERVFRD-1.*, were located.

###### Supplementary Figure 16

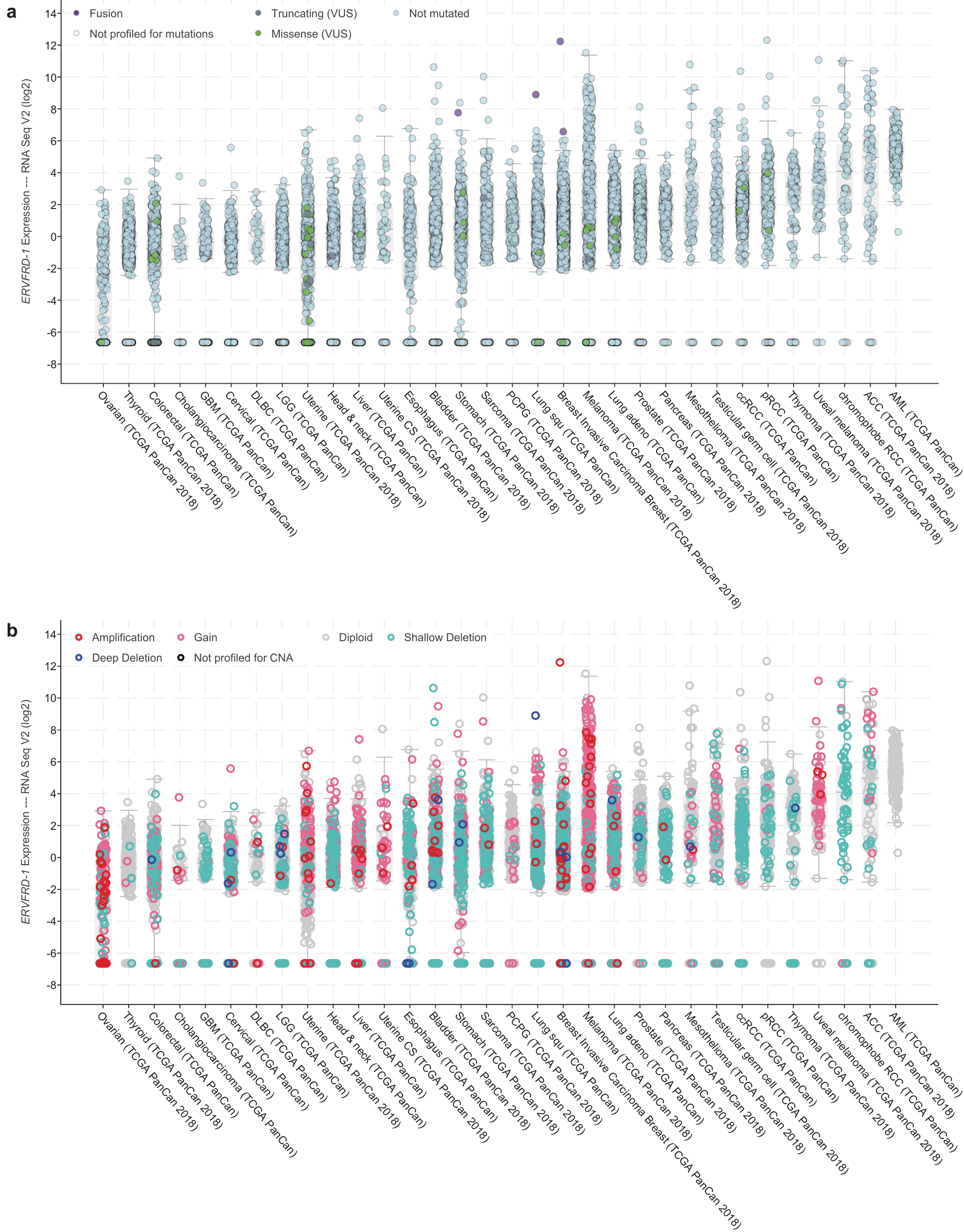

**ERVFRD-1 in TCGA samples.** Expression of *ERVFRD-1* gene across cancer types, colored with somatic mutations and fusions (**a**) and copy number alteration (**b**).

### References

1. Soneson, C., Matthes, K. L., Nowicka, M., Law, C. W. & Robinson, M. D. Isoform prefiltering improves performance of count-based methods for analysis of differential transcript usage. *Genome Biology* **17**, (2016).

2. Saraiva-Agostinho, N. & Barbosa-Morais, N. L. Psichomics: Graphical application for alternative splicing quantification and analysis. *Nucleic Acids Research* **47**, e7 (2019).

3. Falcon, S. & Gentleman, R. Using GOstats to test gene lists for GO term association. *Bioinformatics* **23**, 257–8 (2007).
